# An implantable sensor network for large-scale cellular force mapping in living embryos

**DOI:** 10.64898/2026.10.19.760153

**Authors:** Fei Liu, Jerry D. Wu, Sara Kacmoli, Xiaoyu Zhang, Vanessa Gonzalez, Cullen B. Young, Jinghui Wang, Mengyang Jacky Li, Yuqing Qiu, Gabriella H. Myslo, Angeline Yan, Zhaoyang Song, Yajie Liang, Rebecca D. Burdine, Howard A. Stone, Tian-Ming Fu

## Abstract

Mechanical forces generated by cells are fundamental regulators of embryonic development, yet how locally generated forces are integrated across tissues to drive large-scale morphogenesis remains poorly understood. Addressing this question requires direct quantitative mapping of intercellular forces across tissue scale with cellular resolution while preserving native tissue architectures and dynamics, a capability not yet achieved. Here, we introduce FORCENET, an implantable force-sensing network that enables quantitative tissue-scale force mapping at cellular resolution within living embryos. FORCENET consists of a microfabricated network of ultrathin elastic sensors whose three-dimensional deformations are tracked over time and converted into intercellular force maps through physics-based reconstruction. FORCENET covers hundreds of cells while occupying a total volume 6- to 20-fold smaller than that of a single cell. Anatomical measurements, three-dimensional fluorescent imaging, and cell density and speed tracking confirmed that FORCENET seamlessly integrates within developing embryos without perturbing cell motion and proliferation, tissue architecture, and embryonic development. Applied to zebrafish gastrulation, FORCENET revealed that intercellular forces are dynamically coordinated across supracellular length scales of ∼30–210 μm. Inhibition of fibroblast growth factor signaling accelerated force accumulation and shortened the force-coordination length scales despite only modest changes in cell density and kinematics, demonstrating that FORCENET uncovers information not readily inferred from cell tracking alone. FORCENET establishes a scalable platform for *in vivo* force mapping and reveals that cell-generated forces are coordinated into supracellular mechanical patterns that are regulated by developmental signaling.

## Introduction

Mechanical forces are fundamental regulators of diverse biological processes, including embryonic development, wound healing, tissue regeneration, and cancer progression^1–5^. At the cellular scale, actomyosin contractility, cell–cell adhesion, and cell–matrix interactions generate and transmit forces that regulate cell migration and differentiation^6,7^. However, how these locally generated forces are coordinated to drive tissue-scale dynamics remains a key unanswered question. *In vitro* and *ex vivo* studies have shown that cellular forces can be integrated over supracellular length scales via cell–cell contacts and ECM–mediated mechanical coupling^8–11^. However, because these studies were primarily performed in reconstituted or simplified two-dimensional (2D) systems, whether these observations can be generalized to 3D intact organisms remain unclear. *In vivo* studies have revealed multicellular actomyosin assemblies, coordinated tissue flows, and long-range mechanical feedback^12–14^. However, many of these findings relied on indirect inference of mechanical organizations from structural imaging, cell morphology, or cell kinematics, requiring assumptions about tissue material properties, force balance, and boundary conditions that are often difficult to verify within living organisms^15–17^. As a result, a major gap remains in directly measuring how forces are spatially coordinated and dynamically remodeled within living organisms.

This gap reflects a broader limitation in biological research: whereas gene expression and molecular signals can now be studied with high precision, mechanical forces remain difficult to measure within intact 3D organisms^18–21^. Because mechanical forces *in vivo* are inherently multiscale^22,23^, quantifying their spatial coordination and dynamic remodeling requires direct force mapping across tissue-scale at cellular resolution while preserving native tissue architectures and dynamics^5,24^. Existing technologies fall short of meeting these requirements^18–21^. Single-point probing methods, including micropipette aspiration^25^, laser ablation^12^, and optical tweezers^26^, provide important information, but are spatially localized, low-throughput, or perturbative. Molecular tension sensors^27,28^ and computational force-inference methods^29,30^ offer broader spatial coverage, but their quantitative interpretation depends on calibration or mechanical assumptions that are often difficult to validate *in vivo*. Emerging deformable probes and photonic sensors, including microdroplets^31^, microgels^32^, bioprinted hydrogels^33^, droplet microcavities^34^, deformable microlasers^35^, and upconversion particles^36,37^, provide direct mechanical readout, but remain difficult to deploy densely across tissues for spatially distributed measurements. Therefore, the capability to quantitatively map intercellular forces with tissue-scale coverage, cellular resolution, and minimal perturbation within 3D intact organisms has not yet been achieved.

Here, we present Flexible and Optically Resolvable Cellular forcE-mapping with a Network of Elastic Transducers (FORCENET), an implantable force-sensing network for mapping intercellular forces across living 3D embryos with cellular-scale resolution. Using a monolithically integrated insertion needle, FORCENET can be delivered into targeted embryonic tissues. Survival rate analysis, anatomical measurements, 3D time-lapse imaging, and cell density and motility tracking confirmed that FORCENET does not measurably perturbing cell migration, mitosis, or embryonic development. Endogenous intercellular forces deform individual force-sensing beam elements of FORCENET in 3D. These deformations are captured by 3D fluorescence imaging and converted into quantitative maps of intercellular stress (force per unit beam surface area) through physics-based reconstruction. Applying FORCENET to zebrafish gastrulation reveals spatially heterogeneous and dynamically evolving intercellular stress landscapes organized over supracellular length scales. By integrating these measurements with simultaneous cell tracking, we show that these stress landscapes are more closely associated with local cell density than with instantaneous cell kinematics. Finally, perturbation of fibroblast growth factor (FGF) signaling alters stress accumulation and reduces supracellular length scales despite modest changes in cell density and motion. Together, these results establish FORCENET as a minimally invasive approach for spatiotemporal mapping of intercellular stress and reveal supracellular stress organization as a dynamically regulated feature of embryonic morphogenesis.

## Results

### Concept, design, and characterization of FORCENET

To map intercellular stresses in living embryos, an ideal platform should provide tissue-scale coverage, resolve stresses at the cellular resolution, and integrate within native 3D tissues with minimal footprint. To meet these requirements, we envisioned FORCENET as an ultrathin, fluorescently labeled, elastic polymer network, featuring (i) a force-sensing region spanning across living embryonic tissues, (ii) sensing elements that mechanically couple to surrounding cells and deform in response to endogenous intercellular stresses, and (iii) a total device volume smaller than that of a single cell to minimize disruption to cell–cell adhesion and embryonic development (Fig. 1a). After implantation, the same endogenous intercellular stresses that drive morphogenesis deform FORCENET in 3D, enabling quantitative reconstruction of the stress fields based on optically measured deformation and defined mechanical properties of the sensor (Fig. 1a).

**Fig. 1.**
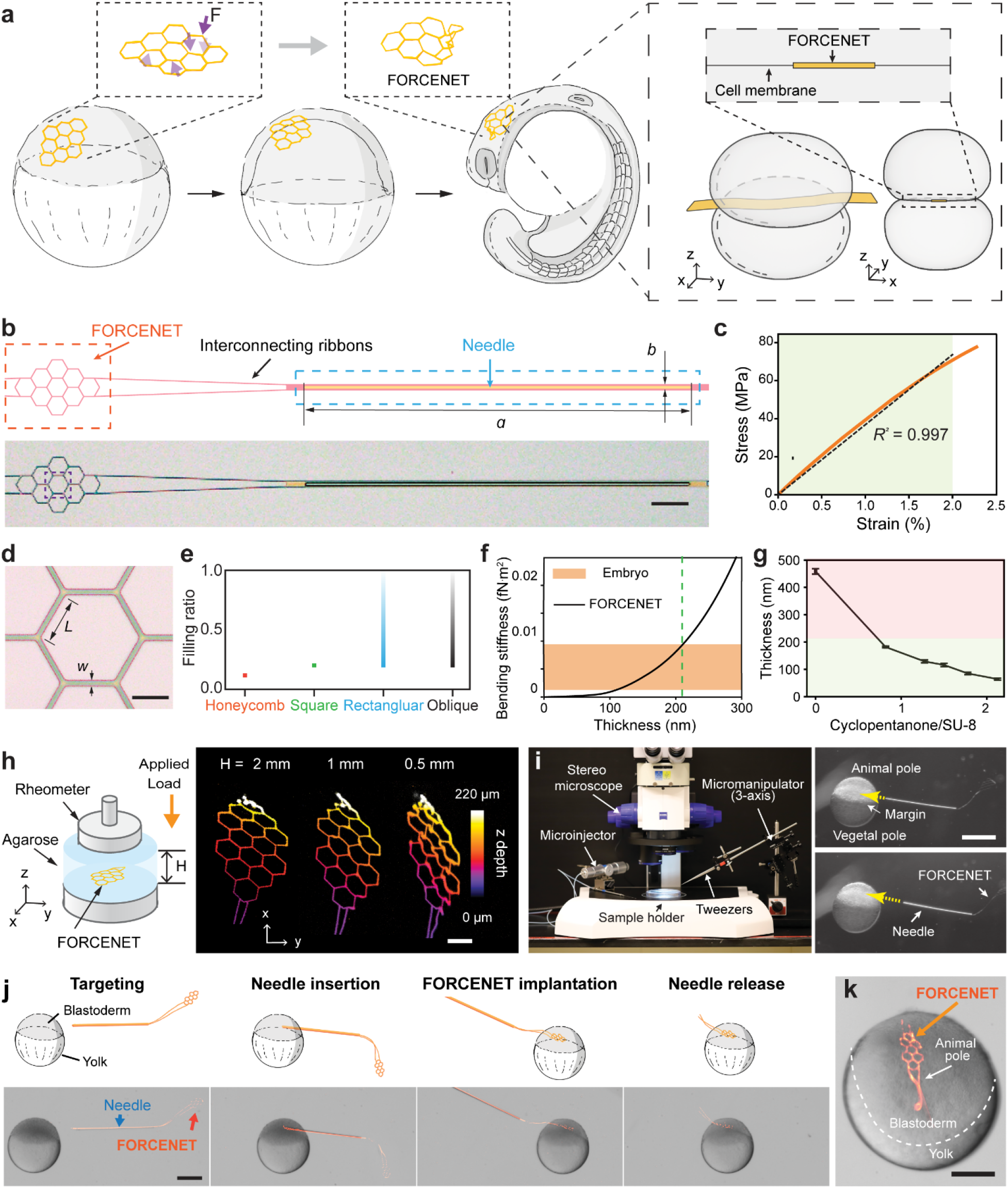
Concept, design, characterization, and implantation of FORCENET for *in vivo* stress mapping. **a,** Schematic of the FORCENET concept. FORCENET is implanted into the zebrafish blastoderm and deforms in response to forces generated during morphogenesis. 3D device deformations are converted into intercellular stresses using mechanical modeling. **Inset:** schematic illustrating the ultrathin cross section of FORCENET (yellow), enabling implantation between cells (gray) while preserving native cell–cell contacts and force transmission. **b,** Device design (top) and corresponding bright-field image (bottom). The device consists of a force-sensing honeycomb lattice region (FORCENET; red dashed box) connected to a stiff implantation needle (blue dashed box) through flexible interconnecting ribbons. *a*, needle length; *b*, needle width. Scale bar, 100 μm. **c,** Representative stress–strain curve obtained from tensile testing of SU-8. The green shaded region indicates the linear strain range used to determine the elastic modulus. Dashed line, linear fit (R^2^=0.997). **d,** Bright-field image of honeycomb lattice sensing architecture. *L*, beam length; *w*, beam width. Scale bar, 20 μm. **e,** Honeycomb lattices achieve the lowest filling ratio among different types of 2D lattices. **f,** Calculated bending stiffness of FORCENET as a function of beam thickness (Supplementary Note 2). Orange shading denotes the estimated stiffness range of a single-cell-thick embryonic tissue layer; the green dashed line indicates the maximum thickness yielding comparable bending stiffness. **g,** Measured beam thickness as a function of cyclopentanone/SU-8 dilution ratio. Green shading indicates the target thickness range in which the beam bending stiffness is comparable to that of a single-cell-thick embryonic tissue layer, whereas red shading indicates thicknesses exceeding this tissue-matched stiffness range. Error bars indicate standard deviation (s.d.). **h,** Schematic of mechanical characterization (left) and progressive deformation of FORCENET under increasing compressive strain captured by 3D imaging (right). FORCENET embedded within agarose was subjected to controlled compressive loading along the z axis. H denotes agarose thickness. Color indicates projection depth. Scale bar, 50 μm. **i,** Custom implantation platform (left) comprising a stereomicroscope, 3-axis micromanipulator, microinjector, and tweezers and representative images demonstrating targeted implantation of FORCENET into the blastoderm (right). Dashed arrows indicate two example insertion trajectories. Scale bar, 500 μm. **j,** Implantation procedure. The needle is aligned with the blastoderm, inserted through the tissue, and subsequently removed after release of the implanted sensing lattice. Scale bar, 300 μm. **k,** Representative embryo following implantation, showing a flat, conformal configuration of FORCENET within the blastoderm. Scale bar, 200 μm.

To realize this concept, we designed the overall device as a monolithically integrated two-part construct consisting of a force-sensing network (FORCENET) and an insertion needle connected by flexible interconnecting ribbons (Fig. 1b). The stiff insertion needle provides the mechanical rigidity needed for targeted delivery into embryonic tissues, whereas the implanted FORCENET remains sufficiently flexible to deform with the surrounding tissues. We chose SU-8 as the material for FORCENET, the insertion needle, and interconnecting ribbons because it enables fabrication of well-defined microscale structures and offers chemical stability and biocompatibility^38^. SU-8 also exhibits linear elastic behavior up to ∼2% strain with a Young’s modulus of ∼4 GPa (Fig. 1c; Extended Data Fig. 1a; Methods), providing a mechanically well-defined framework.

A key design requirement is to maximize spatial coverage of the sensing area while minimizing the amount of material embedded in tissues. Because the material occupancy of a beam network scales with the total beam length for a fixed beam cross-section, we arranged the force-sensing beam elements into a honeycomb lattice (Fig. 1d), inspired by the Honeycomb theorem showing that a regular hexagonal tiling minimizes total perimeter among equal-area partitions of a plane^39^. In specific, for equivalent sensing areas, the honeycomb lattice occupies approximately half or less of the material volume required by square, rectangular, or oblique lattices (Fig. 1e; Extended Data Fig. 1b,c; Supplementary Note 1). In addition, the six-fold rotational symmetry of the honeycomb lattice maximizes angular sampling isotropy among 2D lattices, reducing directional bias in force reconstruction. Furthermore, FORCENET’s macroporous architecture was designed to minimize hindrance of molecular diffusion and cellular movement during embryonic development.

We next selected the dimensions of individual force-sensing beam elements to balance spatial resolution and force sensitivity with minimal tissue perturbation. Each beam element was designed to be 30 µm long and 3 µm wide (Fig. 1d), with a length comparable to typical cell dimensions and a width an order of magnitude smaller to minimize obstruction of cell–cell contacts. The element thickness (∼180 nm) was determined by three competing constraints. First, the elements must be sufficiently compliant to deform under endogenous intercellular stresses while minimizing their mechanical influence on surrounding tissues. Because bending stiffness scales as *Ewh^3^*, where *E*, *w*, and *h* are the Young’s modulus, beam width, and thickness, respectively, this requirement constrains the thickness to be less than ∼210 nm (Fig. 1f; Supplementary Note 2). Second, the resulting deformations must be sufficiently large to be resolved by optical microscopy under physiological stress levels, further limiting the thickness to below ∼200 nm (Extended Data Fig. 1d,e). Third, the elements must remain within the elastic regime under intercellular stresses that can reach the hundreds of pascals to kilopascals range during embryogenesis^31,32^, imposing a lower bound of ∼150 nm (Extended Data Fig. 1f). Together, these considerations define an optimal beam thickness range of 150–200 nm. With this geometry, FORCENET achieves a sensing area of ∼40,000 µm², spanning approximately 100–400 cells, while maintaining a total device volume 6–20-fold smaller than that of a single cell (Extended Data Fig. 1g).

FORCENET was fabricated at high throughput with near-100% yield using standard photolithography (Extended Data Fig. 2a; Methods). Because commercially available SU-8 typically yields films thicker than 500 nm, we diluted SU-8 with cyclopentanone, its primary solvent, to tune film thickness down to ∼60 nm (Fig. 1g). This thickness control enables the force sensitivity and dynamic range of FORCENET to be tailored to different biological systems. Diluted SU-8 was spin-coated onto a silicon wafer coated with a 50-nm-thick sacrificial nickel layer. The thin SU-8 layer was then photolithographically patterned into the FORCENET and the interconnecting ribbons with a thickness of ∼180 nm. A second 10-µm-thick SU-8 layer was then patterned to form the stiff insertion needle. Finally, the sacrificial nickel layer was etched to release the free-standing device. This microfabrication strategy provides substantial flexibility in device geometry, configuration of the force-sensing elements, and beam dimensions (Extended Data Fig. 2b,c), allowing FORCENET to be adapted to different force regimes, developmental stages, and model organisms. Wafer-scale fabrication yields more than 2,000 devices per wafer with uniform film thickness distributions of 182 ± 2 nm (mean ± standard deviation (s.d.)), as confirmed by profilometry measurements (Extended Data Fig. 2d–f).

To enable time-resolved force mapping in living embryos, FORCENET must be optically visible, mechanically stable under physiological conditions, and capable of producing quantifiable 3D deformations under external loads. Because native SU-8 is transparent and exhibits weak autofluorescence, direct visualization of the FORCENET structures is challenging. To achieve high signal-to-noise-ratio (SNR) imaging, we incorporated biocompatible fluorescent dyes into the diluted SU-8, producing stable fluorescence for weeks in E3 medium at 28.5 °C, the standard condition for zebrafish development (Extended Data Fig. 2g–i; Methods). This strategy is compatible with diverse fluorescent dyes spanning the visible spectrum, enabling multiplexed imaging of FORCENET together with fluorescently labeled biological structures in separate spectral channels (Extended Data Fig. 2g). FORCENET can also be visualized under two-photon excitation (Extended Data Fig. 2g), supporting imaging in deeper or scattering tissues. To assess mechanical stability, we measured the Young’s modulus of SU-8 before and after incubation in E3 medium at 28.5 °C and observed no statistically significant change after 7 days (Extended Data Fig. 2j). Finally, we tested whether FORCENET produces detectable 3D deformations in response to controlled external loading. FORCENET was embedded within agarose hydrogels and imaged using an integrated rheometer–spinning-disk confocal setup. Increasing external loads produced corresponding increases in 3D deformation of the force-sensing network (Fig. 1h).

### Targeted implantation of FORCENET into living embryos

The same ultrathin and highly flexible architecture that minimizes tissue perturbation and enhances force sensitivity also creates a major implantation challenge: FORCENET is too compliant to directly penetrate embryonic tissues. To decouple the mechanical requirements for implantation and force sensing, we monolithically integrated a stiff insertion needle with the force-sensing network through flexible interconnecting ribbons (Fig. 1b). The insertion needle, measuring 1.1 mm in length, 8 µm in width, and 10 µm in thickness, has a bending stiffness five orders of magnitude greater than that of the force-sensing network (Supplementary Note 3), providing sufficient rigidity to penetrate the blastoderm.

To enable spatially targeted delivery, we developed a custom implantation platform consisting of a stereomicroscope for real-time visualization and a tweezer mounted on a three-axis micromanipulator to grasp and precisely manipulate the insertion needle (Fig. 1i). This setup allowed us to control the needle entry point and insertion trajectory within optically accessible regions of the blastoderm, enabling FORCENET to be placed into targeted embryonic regions (Fig. 1i; Methods). After insertion, the interconnecting ribbons were severed and the insertion needle was removed, leaving the force-sensing network embedded within the targeted embryonic tissue (Fig. 1j). Widefield fluorescence imaging confirmed that FORCENET remained embedded within embryos without detectable folding or fracture shortly after implantation (Fig. 1k).

### FORCENET integrates with living embryos without measurably perturbing development

A central requirement for *in vivo* force mapping is that the implanted sensor mechanically couples to the surrounding tissue without perturbing embryonic development. We therefore evaluated the FORCENET–embryo interface across organismal, tissue, and cellular length scales. To assess developmental progression at the whole-embryo level, we implanted FORCENET into the blastoderm of zebrafish embryos at ∼4 hours post fertilization (hpf) and initiated correlated bright-field and fluorescence time-lapse imaging at ∼5 hpf (Fig. 2a). Implanted embryos underwent normal development, progressed through the gastrula, completed epiboly by ∼10.4 hpf, and subsequently formed somites without detectable morphological abnormalities (Fig. 2a; Extended Data Fig. 3a; Supplementary Video 1). During gastrulation, large-scale tissue deformation at the animal pole induced bending of the FORCENET beam elements, indicating mechanical coupling between FORCENET and surrounding embryonic tissues.

**Fig. 2.**
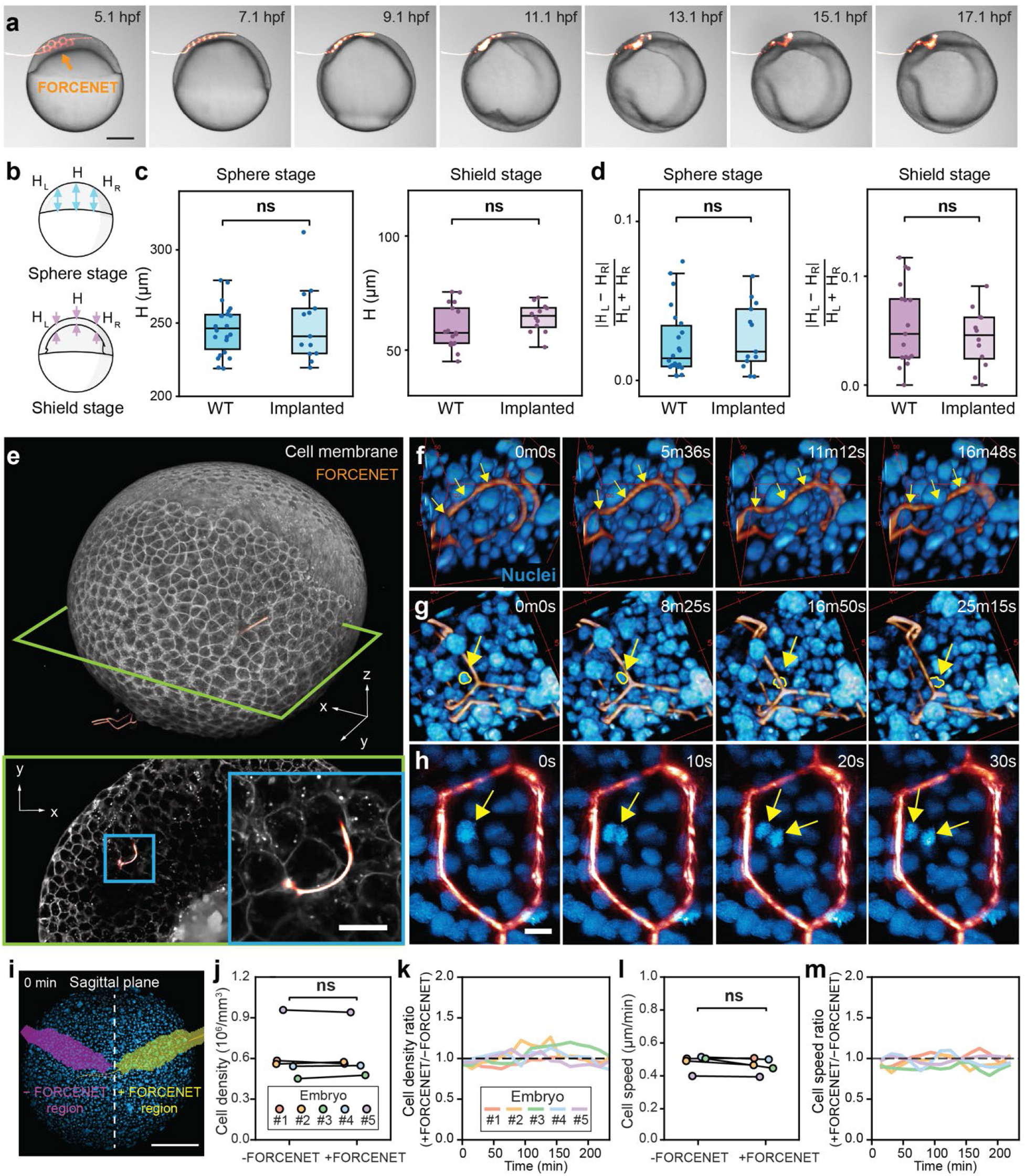
FORCENET integrates with living embryos without measurably perturbing development. **a,** Correlative brightfield-fluorescence time-lapse imaging of zebrafish development following FORCENET (orange) implantation (see also Supplementary Video 1). Scale bar, 200 μm. **b,** Schematic defining embryo morphology measurements at the sphere and shield stages. H denotes blastoderm height at the embryonic midline; H_L_ and H_R_ denote lateral heights. **c,** Quantification of blastoderm height in non-implanted wildtype (WT) control and implanted embryos at the sphere and shield stages. Box plots show the median (center line), interquartile range (box), and range of non-outlier values (whiskers), with individual embryos overlaid as dots. At the sphere stage, n = 22 for WT and n = 13 for implanted embryos; at the shield stage, n = 17 for WT and n = 12 for implanted embryos. Statistical significance was assessed using a two-tailed unpaired t-test. No significant difference was observed at either stage (sphere stage, p = 0.694; shield stage, p = 0.166; ns, not significant). **d,** Quantification of embryo asymmetry. Asymmetry is defined as the normalized difference between H_L_ and H_R_. Statistical significance was assessed using the Mann–Whitney U test. No significant difference was observed at either stage (sphere stage, p = 0.597; shield stage, p = 0.479). **e,** 3D two-photon imaging of the interface between FORCENET (orange) and surrounding cell membranes (gray) (see also Supplementary Video 2). Top, 3D rendering; bottom, corresponding 2D cross-sectional slice. The green plane indicates the slice position and the blue box highlights the magnified region shown in the zoom-in view. Scale bar, 20 μm. **f,** 3D two-photon imaging showing dynamic deformation of a FORCENET beam by neighboring cells (yellow arrows; see also Supplementary Video 3). Bounding box dimensions: 84 × 84 × 82 μm^3^. **g,** 3D two-photon imaging showing a cell (yellow arrows) migrating through the macropores of FORCENET (see also Supplementary Video 3). Bounding box dimensions: 92 × 97 × 90 μm^3^. **h,** Maximum-intensity projection (MIP) of lattice light-sheet with adaptive optics imaging showing cell mitosis (yellow arrows) adjacent to FORCENET (see also Supplementary Video 3). Scale bar, 10 μm. **i,** Definition of the implanted region and its contralateral counterpart. The device segmentation of FORCENET has been dilated to encompass the region near the device (+FORCENET, yellow) and reflected across an anatomical plane of symmetry to create a comparison volume without the device (−FORCENET, pink). Scale bar: 200 μm. **j,** Comparison of cell density between +FORCENET and −FORCENET regions. Each pair of connected points represents one embryo. No statistically significant difference was observed (two-tailed paired t-test; n = 5; p = 0.903). **k,** Temporal evolution of the cell-density ratio between +FORCENET and −FORCENET regions. Colors indicate individual embryos. **l,** Comparison of mean cell speed between +FORCENET and −FORCENET regions. No significant difference was observed (p = 0.131). **m,** Temporal evolution of the cell speed ratio between +FORCENET and −FORCENET regions.

To further assess developmental compatibility, we measured survival and morphology at 24 hpf in a larger cohort of implanted embryos (n = 12) and non-implanted controls (WT, n = 29). All implanted embryos survived to 24 hpf, and more than 80% developed without detectable morphological abnormalities (Extended Data Fig. 3b). We next compared key morphometric features between normally developing implanted embryos and control embryos at the sphere and shield stages (Fig. 2b–d; Methods). Measurements of blastoderm thickness and embryo asymmetry revealed no statistically significant differences between the two groups at either developmental stage (Fig. 2c,d). Together, these organismal-scale analyses indicate that FORCENET implantation does not measurably perturb embryonic morphogenesis.

We next examined local tissue integration using high-resolution 3D time-lapse imaging. Two-photon imaging of the FORCENET–cell membrane interface revealed no visible cavities, separations, or tissue discontinuities surrounding the implanted FORCENET (Fig. 2e; Supplementary Video 2), indicating close integration of FORCENET within the blastoderm. This tight tissue contact is essential for transducing endogenous intercellular stresses to the force-sensing beam elements. Moreover, time-lapse two-photon imaging of implanted embryos with endogenously labeled nuclei showed that surrounding cells dynamically deformed the force-sensing beam elements during development (Fig. 2f; Supplementary Video 3), demonstrating that FORCENET is sufficiently compliant to respond to endogenous intercellular stresses *in vivo*. Furthermore, cells were observed to migrate through the macroporous architecture of FORCENET (Fig. 2g; Supplementary Video 3), indicating that the network does not physically obstruct tissue remodeling or cell movements.

To investigate whether FORCENET perturbs cellular dynamics at higher spatiotemporal resolution, we performed rapid 3D imaging using lattice light-sheet microscopy with adaptive optics^40^. We observed complete mitotic events adjacent to implanted FORCENET (Fig. 2h; Supplementary Video 3), suggesting that cells near FORCENET retain normal proliferative dynamics.

Finally, we quantitatively compared local cell density and motility between implanted regions (+FORCENET) and contralateral control regions (−FORCENET) within the same embryos (Fig. 2i, Extended Data Fig. 4a,b; Methods). Across the 250-min imaging session, cell density and speed showed no statistically significant differences between the +FORCENET and −FORCENET regions (Fig. 2j–m; Extended Data Fig. 4c–e). Collectively, these results demonstrate that FORCENET integrates mechanically with developing embryonic tissues while preserving whole-embryo morphogenesis, local tissue architecture, as well as cell migration and proliferation.

### Spatiotemporal stress mapping within developing zebrafish embryos

Having established that FORCENET integrates with living embryos without measurably perturbing development, we next used it to map intercellular stress dynamics during zebrafish gastrulation, when cell-generated forces are transmitted through dynamic cell–cell and cell–matrix interactions to sculpt tissue-scale morphogenesis^41,42^. A representative implanted embryo at ∼6 hpf showed close integration of FORCENET within the blastoderm, with force-sensing beam elements maintaining tight contact with surrounding cells (Fig. 3a; Extended Data Fig. 5). Different beam elements exhibited distinct 3D deformations (Extended Data Fig. 5b), confirming that FORCENET can resolve spatially heterogeneous mechanical activity within developing embryonic tissues.

**Fig. 3.**
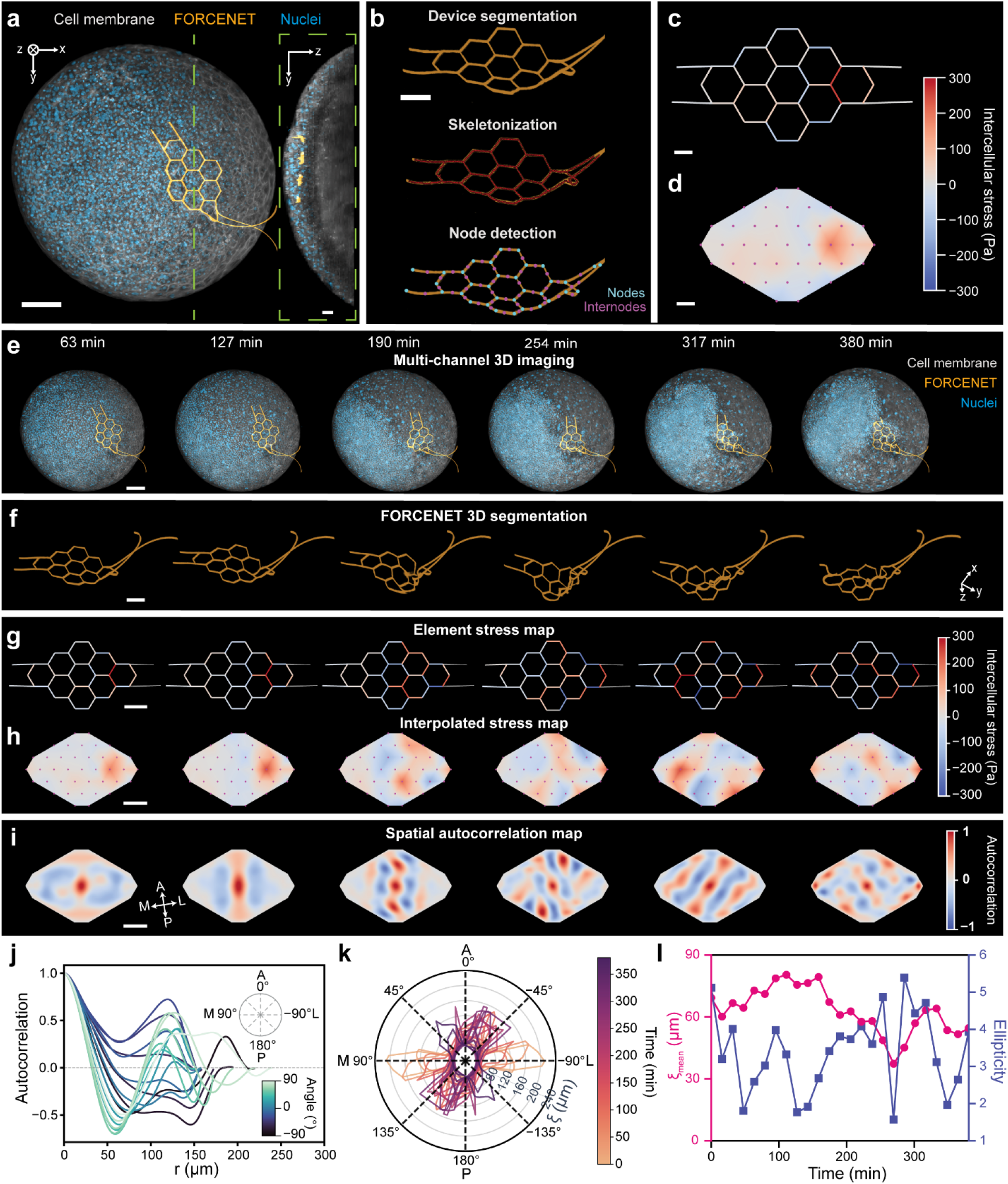
FORCENET maps spatiotemporal intercellular stress dynamics during gastrulation. **a,** Representative 3D rendering of an implanted embryo showing cell membranes (gray), nuclei (blue), and FORCENET (orange). Inset, yz-view MIP of 20 optical slices highlighting FORCENET embedded within the blastoderm. **b,** Representative images of the three major computer-vision tasks used to extract FORCENET geometry from raw two-photon images: device segmentation, skeletonization of the segmented device, and detection of nodes and internodes from the skeletons (see also Supplementary Video 4). **c,** Intercellular stress mapped onto the beam elements. **d,** Interpolated intercellular stress field. Panels **a**–**d** show a representative time point, t = 0 min (∼6 hpf). **e,** 3D renderings of the same implanted developing embryo over time (see also Supplementary Video 5). **f,** Corresponding segmented FORCENET. **g,** Element-wise intercellular stress maps. **h,** Interpolated intercellular stress fields. **i,** Normalized spatial autocorrelation maps of intercellular stress. **j,** Orientation-dependent spatial autocorrelation profiles of intercellular stress at t = 317 min (∼11.3 hpf). Curves correspond to different angular directions relative to the embryonic axes (inset). **k,** Polar representation of stress correlation length, *ξ*, as a function of orientation and developmental time. Correlation lengths were determined from the first zero crossing of the autocorrelation profiles. **l,** Mean correlation length *ξ*_mean_ (magenta) and ellipticity (blue, calculated from *ξ*_max_/*ξ*_min_) are plotted as functions of time. Scale bars, 100 μm (**a** main, **e**, **i**), 50 μm (**b**, **g**–**h**), 30 μm (**c**, **d**), and 25 μm (**a** inset). In (**i**–**k**), A, anterior; P, posterior; M, medial; L, lateral.

To convert FORCENET deformation into quantitative stress maps, we developed an image-processing and physics-based reconstruction pipeline (Fig. 3b; Extended Data Fig. 6a; Supplementary Video 4; Methods). FORCENET was first segmented from raw 3D fluorescence image stacks and skeletonized to extract the network geometry. Second, node positions, corresponding to vertices of the honeycomb network, were identified and used to calculate nodal translations relative to the undeformed configuration (Extended Data Fig. 6b; Supplementary Video 4). In parallel, nodal rotations were estimated by tracking internodal positions along individual beam elements (Extended Data Fig. 6c; Supplementary Video 4; Methods). Third, these measured translations and rotations were used as inputs for finite element analysis (FEA) to obtain nodal forces, which are then converted to effective distributed loads and subsequently to stresses on individual beam elements (Fig. 3c; Extended Data Fig. 6a; Methods). An uncertainty estimation incorporating the two dominant error sources—beam thickness variation and nodal localization uncertainty—yielded a relative stress-sensing uncertainty of ∼7.6% (Supplementary Note 4). Last, the stresses on individual beam elements were spatially interpolated to characterize intercellular stress distribution across the covered tissue regions (Fig. 3d; Methods). Characterization using stress gratings with varying spatial length scales and orientations confirmed that the reconstruction pipeline preserves stress spatial distributions over length scales comparable to or larger than the beam element length, while higher spatial frequency stress variations are averaged by the finite sampling of FORCENET (Extended Data Fig. 7). The reconstructed stress maps revealed spatially heterogeneous intercellular stress distributions across the blastoderm, showing mean signed stress of 11.8 Pa ± 73.8 Pa (mean ± s.d.) and mean absolute stress of 72.5 Pa ± 74.3 Pa (mean ± s.d.) across the representative sensing region (Fig. 3d).

We next applied this pipeline to time-lapse measurement data beginning at ∼6 hpf to track the spatiotemporal evolution of the intercellular stress field during gastrulation (Fig. 3e–h; Extended Data Fig. 5a; Supplementary Video 5). Multi-channel time-lapse imaging revealed progressive deformation of FORCENET as large-scale morphogenetic movements reshaped the blastoderm (Fig. 3e). Segmentation of the FORCENET channel enabled direct visualization of its evolving 3D network geometry over time (Fig. 3f). Initially, FORCENET remained relatively flat. As gastrulation progressed, spatially heterogeneous deformations emerged and propagated across the sensing region (Fig. 3f; Extended Data Fig. 5a). Using these measured deformations, we obtained dynamic intercellular stress fields and visualized them both on the FORCENET honeycomb lattice (Fig. 3g) and as interpolated stress maps (Fig. 3h). The reconstructed intercellular stress landscapes exhibited marked dynamic reorganization throughout gastrulation: as gastrulation progressed, domains of elevated stress emerged, shifted, and changed sizes over time (Fig. 3g,h; Extended Data Fig. 5a; Supplementary Video 5). Notably, regions of peak stress appeared sequentially along the mediolateral axis of the embryo (Extended Data Fig. 5c,d; Supplementary Video 5), suggesting coordinated propagation of mechanical activity across the developing tissue. Together, these measurements reveal that gastrulation is accompanied by spatially heterogeneous and dynamically evolving intercellular stress landscapes.

We next asked whether these intercellular stress landscapes were spatially organized over supracellular scales. To quantify the correlation length scales of intercellular stress, we calculated the spatial autocorrelation of the reconstructed stress field^8^ (Fig. 3i; Supplementary Note 6; Methods). The resulting autocorrelation maps contained alternating positive and negative correlation domains (Fig. 3i), indicating that intercellular stresses were spatially organized over characteristic length scales. In addition, these domains were anisotropic, suggesting direction-dependent mechanical coordination within the gastrulating tissue (Fig. 3i). Furthermore, the spatial autocorrelation maps evolved from relatively simple patterns into increasingly complex and oscillatory structures over time (Fig. 3i), indicating progressive remodeling of tissue-scale mechanical organization during morphogenesis. To quantify directional dependence, we extracted radial profiles at a representative time point with orientation defined relative to the embryonic anatomical axes (Fig. 3j). These profiles showed pronounced directional differences in decay behavior and oscillatory structure: profiles along the medial direction exhibited stronger oscillations with alternating positive and negative correlation domains, whereas profiles along the lateral direction showed weaker oscillations and more monotonic decay (Fig. 3j). We defined the stress correlation length, *ξ*, as the first zero-crossing distance of each radial autocorrelation profile when applicable. At this representative time point, *ξ* ranged from ∼30 µm at 53° to ∼150 µm at −50° relative to the anterior direction, demonstrating anisotropic supracellular stress coordination.

We then asked how this supracellular organization changes during gastrulation by extending the radial autocorrelation analysis across the full 380-min time series (∼6-12 hpf, Fig. 3k). The angular distributions of *ξ* revealed anisotropic and dynamically evolving stress coordination during gastrulation. The maximum stress correlation length, *ξ*_max_, reached ∼210 µm near the onset of the imaging session along the mediolateral axis (∼89.7° from the anterior direction), indicating long-range coordination of intercellular stresses. To further characterize the temporal remodeling of supracellular organization, we quantified the mean stress correlation length, *ξ*_mean,_ which reports the average length scale of stress coordination, and the ellipticity of the autocorrelation fields, which measures the degree of anisotropy in the stress landscape, over time (Fig. 3l; Methods). *ξ*_mean_ changed gradually over the course of gastrulation, whereas autocorrelation ellipticity fluctuated on a faster time scale, suggesting that the range and orientation of tissue-scale mechanical coordination are regulated by distinct dynamic processes. Together, these analyses provide a quantitative measurement of supracellular stress length scales and reveal their continuous remodeling during embryonic morphogenesis.

Finally, we asked whether the intercellular stress accumulation, spatial heterogeneity, and dynamic anisotropy observed in the representative embryo were reproducible across embryos. To address this, we examined intercellular stress dynamics across a cohort of six gastrulating embryos (Extended Data Fig. 8a, b). We observed that the mean intercellular stress averaged over all force-sensing elements progressively increased during gastrulation (Extended Data Fig. 8c; Supplementary Video 6). In parallel, the standard deviation of the stress fields also increased over time (Extended Data Fig. 8d), indicating growing spatial variability in stress magnitude as morphogenesis progressed. By contrast, the autocorrelation ellipticity fluctuated dynamically without exhibiting a clear global trend (Extended Data Fig. 8e), consistent with the observed continual reorientation of tissue-scale mechanical anisotropy during gastrulation (Fig. 3k,l). These cohort-level measurements support the observation that embryonic morphogenesis is accompanied by progressive stress accumulation, increasing spatial heterogeneity, and dynamic remodeling of supracellular stress organization.

### Simultaneous intercellular stress mapping and cell tracking

Next, we asked whether intercellular stress landscapes are reflected in commonly measured cellular readouts, such as local cell density and instantaneous cell kinematics. Addressing this question requires simultaneous measurements of intercellular stresses and cellular dynamics within the same region of a developing embryo. We therefore combined FORCENET-based stress mapping with 3D cell segmentation and tracking (Extended Data Fig. 9a–f). We first applied our reconstruction pipeline to map temporally evolving intercellular stress fields from 3D time-lapse imaging of a FORCENET-implanted embryo (Fig. 4a; Extended Data Fig. 9c). Time-resolved quantification showed that both the mean and spatial standard deviation of the intercellular stress field increased progressively during gastrulation, with a more pronounced rise at later developmental stages (Extended Data Fig. 9g). We next performed 3D cell segmentation and tracking from nuclear images within regions surrounding FORCENET using Cellpose-SAM^43^ and Ultrack^44^ (Fig. 4b,c; Extended Data Fig. 9b; Methods). Cell segmentation yielded spatial maps of local cell density (Fig. 4b; Extended Data Fig. 9d), whereas tracking of individual nuclei enabled reconstruction of cell speed and acceleration magnitude fields (Fig. 4c; Extended Data Fig. 9e,f; Supplementary Video 7; Methods). These cellular metrics revealed distinct temporal behaviors: consistent with previous studies^45,46^, spatially averaged cell density gradually increased during early gastrulation before reaching a plateau at ∼300 min after gastrulation onset (Extended Data Fig. 9h). By contrast, mean cell speed and acceleration magnitude remained relatively stable throughout the imaging session, with only modest temporal fluctuations (Extended Data Fig. 9i,j).

**Fig. 4.**
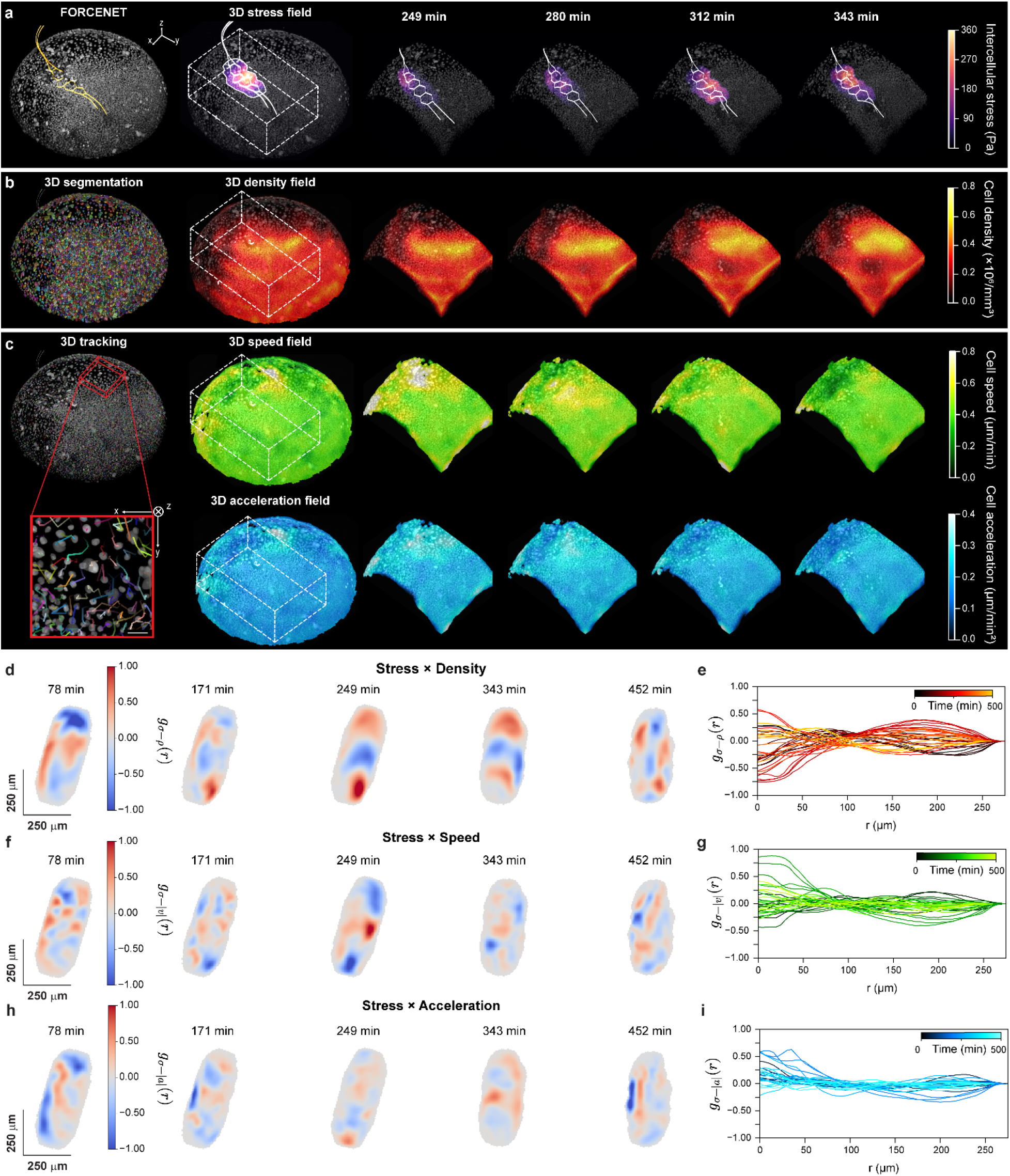
Simultaneous intercellular stress mapping and cell tracking. **a,** Representative 3D stress field measured by FORCENET. Left, 3D imaging of FORCENET (orange) and nuclei (white) within the blastoderm. Middle, reconstructed intercellular stress field. Right, time series of 3D images overlaid with reconstructed intercellular stress fields. The solid outline indicates the region surrounding FORCENET used for stress-field reconstruction. Imaging onset (t = 0 min) is ∼6 hpf. **b,** Nuclear segmentation and reconstructed cell-density field from the same embryo shown in **a**. Left, 3D nuclear segmentation, with colored objects indicating individual segmented nuclei. Middle, reconstructed cell-density field. Right, time series of reconstructed cell-density fields. **c,** Cell tracking and reconstructed cell-speed fields (top) and cell-acceleration magnitude fields (bottom) from the same embryo in **a**. Left, 3D nuclear tracking, with the inset showing representative nuclear trajectories. Middle, reconstructed 3D cell-speed field and cell-acceleration magnitude field. Right, time series of reconstructed cell-speed and cell-acceleration magnitude fields. The same dashed box in **(a–c)** denotes the region analyzed for cell density and kinematics. **(d–i)** Spatial cross-correlation functions between intercellular stress and cell density (**d**, **e)**, cell speed (**f**, **g**), and cell acceleration magnitude (**h**, **i**) throughout gastrulation. (**d**, **f**, **h**) Representative 2D slices of the corresponding 3D cross-correlation maps at t = 78 min, 171 min, 249 min, 343 min, and 452 min. Curves in (**e**, **g**, **i**) are radial averaged cross-correlation functions color-coded by imaging time.

We then quantified how the reconstructed intercellular stress landscapes related spatially to local cell density, speed, and acceleration magnitude. To this end, we computed spatial cross-correlation functions between stress fields and maps of each cellular readout within the same tissue regions throughout gastrulation (Extended Data Fig. 9b; Supplementary Note 6; Methods). Stress-density cross-correlations exhibited distance-dependent structures, with positive and negative domains that evolved over time (Fig. 4d,e). By contrast, cross-correlations between stress and cell speed, as well as between stress and cell acceleration magnitude, were weaker (Fig. 4e), and radial averaged profiles decayed over relatively short distances (Fig. 4f). These results indicate that instantaneous cell kinematics alone incompletely capture the spatial organization of intercellular stresses. Together, these analyses suggest that intercellular stress landscapes are more closely associated with local cell density than with instantaneous cell speed or acceleration magnitude. Moreover, the maximum cross-correlation coefficient between stress and cellular readouts remained moderate, indicating that direct stress mapping reveals mechanical information not readily inferred from cell density or cell motion alone.

### FGF signaling perturbation remodels supracellular stress organization

Finally, we investigated whether developmental signaling pathways regulate supracellular stress organization during gastrulation. We focused on FGF signaling, a key regulator of gastrulation and body-axis formation, and inhibited FGF receptor tyrosine kinases using SU5402 (Fig. 5a; Extended Data Fig. 10a; Methods)^47–49^. Embryos were treated with 10-μM SU5402 beginning at 30% epiboly, whereas control embryos were maintained in E3 medium (Fig. 5b). Consistent with the established role of FGF signaling in gastrulation and body-axis formation^50^, SU5402-treated embryos developed phenotypic abnormalities, including a ∼40% reduction in body length compared with controls at 24 hpf (Fig. 5c; Extended Data Fig. 10b).

**Fig. 5.**
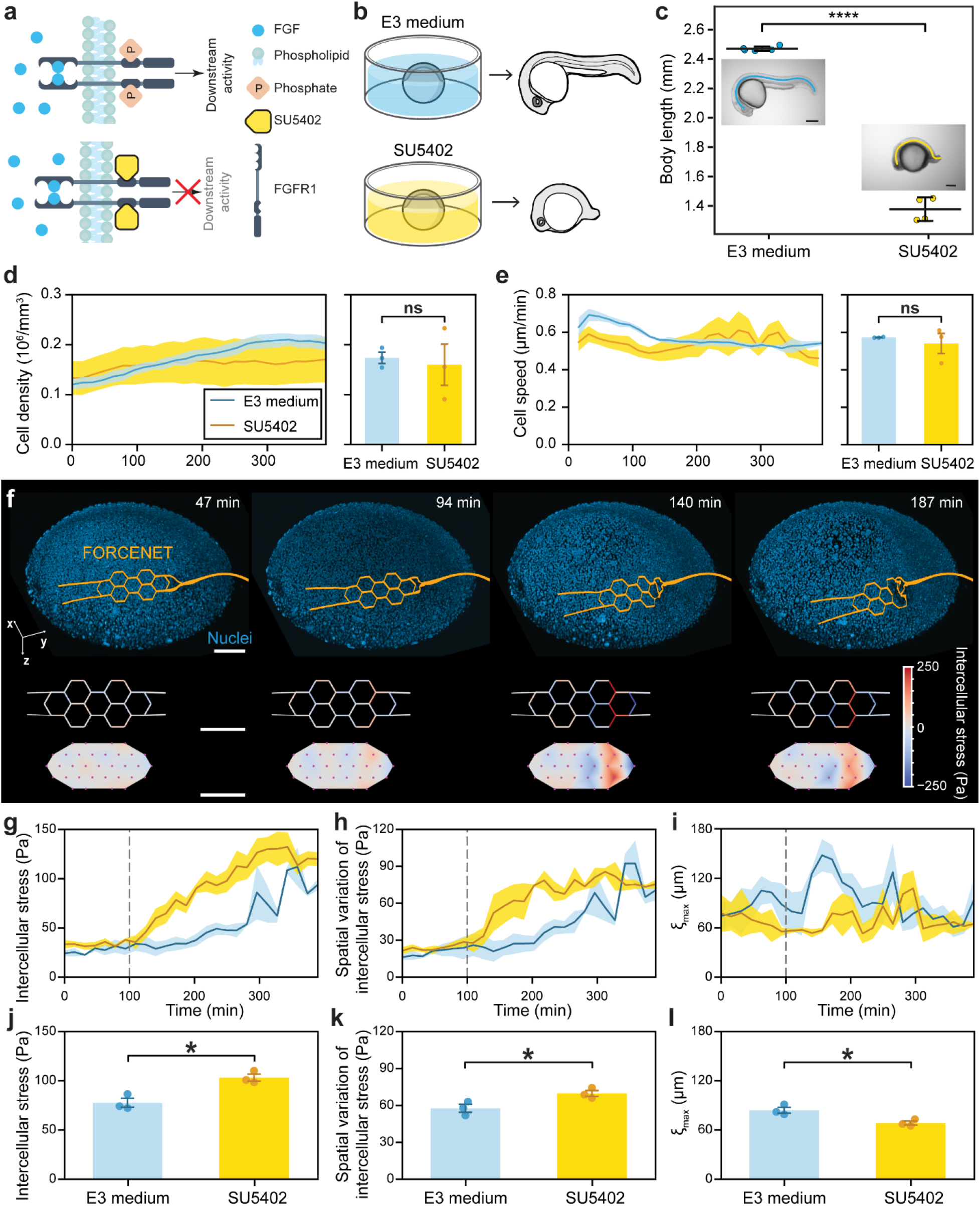
FGF signaling perturbation remodels supracellular stress organization. **a,** Schematic of SU5402 inhibition of FGF signaling by blocking FGF receptor phosphorylation and downstream signaling. **b,** Schematic showing the experimental design for SU5402 treatment and representative morphological outcomes at 24 hpf in control embryos maintained in E3 medium and embryos treated with SU5402. **c,** Quantification of body length at 24 hpf in control (E3 medium) and SU5402-treated embryos. Error bars indicate mean ± s.d.. Statistical significance was determined using a two-tailed Welch’s t-test (n = 5 control embryos and n = 4 SU5402-treated embryos; p = 7.58 × 10^-5^). Insets show representative bright-field images of control and SU5402-treated embryos at 24 hpf. Colored lines indicate the body axis used for length measurement. Scale bar, 200 μm. **(d, e)** Temporal evolution of cell density **(d),** and cell speed **(e)** in control and SU5402-treated embryos. Lines indicate group means and shaded regions denote ±1 standard error of the mean (s.e.m.). Bar plots show the mean time-averaged cell density **(d)** and speed **(e)** across embryos (n = 3 per group), with individual dots representing time-averaged values per embryo, and error bars denoting ± s.e.m. No significant differences were detected between groups based on a two-tailed Welch’s t-test. The p-values for cell density and cell speed were 0.771 and 0.607, respectively. **f,** Representative time-lapse stress mapping in a SU5402-treated embryo. Top, 3D renderings of nuclei (blue) and FORCENET (orange). Middle, reconstructed element-wise intercellular stress maps. Bottom, interpolated intercellular stress fields. Scale bars, 100 μm. **(g–i)** Temporal evolution of mean intercellular stress (**g**), spatial variation of intercellular stress, defined as the standard deviation of stress measurements across FORCENET elements at each time point (**h**), and maximum stress correlation length, *ξ*_max_ (**i**). Solid lines indicate group means and shaded regions denote ± s.e.m. Dashed lines mark the onset of stress accumulation. **(j–l)** Bar plots show the time-averaged means of intercellular stress (**j**), spatial variation of intercellular stress (**k**), and *ξ*_max_ (**l**) for individual embryos (n = 3 per group), with individual dots indicating time-averaged values per embryo, and error bars denoting ± s.e.m. Imaging onset (t = 0 min) for (**d**-**l**) is ∼6 hpf. Time averages were computed from t = 100 min to the end of the imaging window. Statistical significance was determined using a two-tailed Welch’s t-test: p = 0.012 (stress), p = 0.044 (spatial variation), p = 0.029 (*ξ*_max_).

We first asked whether these phenotypic abnormalities were accompanied by detectable changes in local cellular organization or motion. Using 3D cell segmentation and tracking, we quantified local cell density, speed, and acceleration magnitude in control and SU5402-treated embryos (Fig. 5d,e; Extended Data Fig. 10c). Cell density showed broadly similar temporal profiles between the two groups during the first ∼200 min of imaging, with only a modest reduction emerging in SU5402-treated embryos at later stages (Fig. 5d). Similarly, although cell speed and acceleration magnitude were slightly reduced in SU5402-treated embryos during early gastrulation, they substantially overlapped between the two groups throughout the imaging period, and time-averaged measurements did not differ significantly (Fig. 5e; Extended Data Fig. 10d). Thus, within the imaged regions, local cell density and kinematic readouts captured only modest effects of FGF inhibition.

We next asked whether FORCENET could reveal mechanical changes caused by FGF perturbation that were not readily apparent from cellular readouts. We performed time-lapse two-photon imaging beginning ∼2 hours after FORCENET implantation (∼6 hpf) and extracted intercellular stress fields using the reconstruction pipeline (Fig. 5f; Extended Data Fig. 10c). In both control and SU5402-treated embryos, FORCENET progressively deformed during gastrulation, revealing dynamically evolving and spatially heterogeneous stress landscapes (Fig. 5f). However, unlike the cell density and kinematic measurements, reconstructed intercellular stress fields showed marked differences between control and SU5402-treated embryos (Fig. 5g,h). Stress accumulation began earlier in SU5402-treated embryos, at ∼7.6 hpf (∼100 min after imaging onset), compared with ∼9 hpf (∼180 min after imaging onset) in controls, and increased more rapidly over time (Fig. 5g,h). During the first 100 min after imaging onset, control and SU5402-treated embryos showed no significant differences in mean intercellular stress magnitude, stress-field spatial variability, or maximum correlation length (Extended Data Fig. 10e). After ∼100 min, however, SU5402-treated embryos exhibited higher mean intercellular stress magnitude than controls (103.2 ± 6.2 Pa vs. 77.7 ± 7.7 Pa, mean ± s.d.), as well as greater spatial variability in intercellular stresses (69.7 ± 4.2 Pa vs. 57.6 ± 5.6 Pa; Fig. 5j,k).

Last, we asked whether FGF perturbation also altered the supracellular length scale over which intercellular stresses were coordinated. Spatial autocorrelation analysis beginning 100 min after imaging onset showed that the maximum stress correlation length was reduced in SU5402-treated embryos compared with controls (68 ± 4 µm vs. 84 ± 6 µm; Fig. 5i,l), indicating that the elevated stresses induced by FGF signaling disruption were coordinated over shorter supracellular length scales. Therefore, FGF signaling inhibition increased intercellular stress magnitude and spatial variability while reducing supracellular stress-coordination length scales, despite modest changes in local cell density and instantaneous cell kinematics. Together, these results indicate that FGF signaling regulates tissue-scale mechanical organization during gastrulation, extending previous evidence that FGF signaling controls morphogenetic movements^49,51^, adhesion-associated epithelial-to-mesenchymal transition^51^, cytoskeletal remodeling^52^, macroscopic force generation^53^, and tissue mechanics^54^. These findings further demonstrate that FORCENET can uncover mechanical phenotypes not readily inferred from imaging-based measurements of cellular organization and motion alone.

## Discussion

We have developed FORCENET, an implantable force-sensing network that enables minimally invasive mapping of intercellular stress dynamics in living embryos. By integrating an ultrathin macroporous elastic transducer network, a targeted implantation method, high-resolution 3D fluorescence imaging, and physics-based reconstruction, FORCENET maps intercellular stresses across tissue-scale regions while preserving cellular resolution. FORCENET occupies a total volume smaller than that of a single cell, integrates mechanically with surrounding tissues, and does not measurably perturb embryonic morphology, local cell density, cell migration or proliferation. These features establish FORCENET as a scalable platform for longitudinal stress mapping in living developmental systems.

FORCENET addresses a central limitation of existing *in vivo* force-sensing technologies: the trade-off among spatiotemporal coverage, resolution, quantitative force readout, and invasiveness. Existing localized force sensors, including microdroplets^31^, microgels^32^, droplet microcavities^34^, deformable microlasers^35^, and upconversion particles^36,37^, provide important direct force readouts in living tissues, but large-scale mapping requires either dense deployment of many inclusions or larger individual sensors, whose volume scales as the 3/2 power of the projected sensing area. By contrast, FORCENET achieves spatial coverage through an ultrathin honeycomb network whose material volume scales linearly with sensing area, spanning hundreds of microns (∼100–400 cells) while maintaining a total material volume smaller than that of a single cell (Extended Data Fig. 1g). The open macroporous architecture of FORCENET permits cell migration and mitosis through the sensing region, helping preserve native tissue remodeling. Because all force-sensing elements are microfabricated with defined geometric and material properties, measurements across locations and embryos can be interpreted within a common mechanical framework without extensive sensor-by-sensor calibration. Together, these features distinguish FORCENET from localized force sensors and enable spatially distributed stress mapping in dynamic 3D tissues.

Beyond its technological advance, FORCENET reveals supracellular stress organization as a dynamic feature of embryonic morphogenesis. Previous *in vitro*, *ex vivo*, and *in vivo* studies have shown that mechanical activity can be coordinated over multicellular distances^8–14^, yet direct spatially distributed mapping of intercellular stress organization within intact vertebrate embryos has not yet been achieved. By mapping intercellular stress fields during zebrafish gastrulation, we found that stresses are spatially heterogeneous, dynamically remodeled, and coordinated over supracellular length scales spanning 30–210 µm. The correlation length and anisotropy of these stress fields changed over time, demonstrating that supracellular stress organization is continuously remodeled during morphogenesis.

Simultaneous stress mapping and cell tracking further show that intercellular stress measurement contains information not readily inferred from common cellular readouts. Stress landscapes were more closely associated with local cell density than with instantaneous cell speed or acceleration magnitude, yet the correlations remained moderate. Thus, cell density and kinematics provide useful but incomplete proxies for the embryonic mechanical state and its time evolution. FORCENET therefore complements such cellular readouts by providing direct mechanical measurements that relate tissue structure, cell motion, and stress organization within the same developing embryo.

The FGF perturbation experiments further suggest that supracellular stress organization is developmentally regulated. FGF inhibition produced only modest changes in local cell density and instantaneous cell kinematics, but significantly altered stress dynamics: stress accumulation began earlier, stress magnitude and spatial variability increased, and the maximum stress-correlation length decreased. These results show that developmental signaling perturbation can reshape tissue-scale mechanical organization in ways not readily apparent from cellular readouts alone. Together, these findings suggest that intercellular stress can serve as a sensitive readout of developmental perturbation.

Several limitations and opportunities remain. In its current implementation, FORCENET primarily reports stress components associated with deformation along the thickness dimension of the force-sensing beam elements and does not capture the full stress tensor. Future device architectures, such as stretchable materials or designs^55,56^, mechanical metastructures^57^, and nonplanar geometries^58^, could enable reconstruction of additional stress components. In addition, the measurable force range and sensitivity could also be improved through the use of softer materials^56^ and enhanced optical readouts using super-resolution imaging^40^. Moreover, although this study focused on zebrafish gastrulation, FORCENET is intrinsically modular: lattice geometry, beam dimensions, mechanical stiffness, and fluorescent labeling can be systematically tuned for different force ranges, imaging modalities, developmental stages, and model systems. Preliminary experiments showing stable long-term integration in developing human cortical organoids suggest that FORCENET can be extended beyond zebrafish embryos (Supplementary Fig. 1). Furthermore, the microfabrication-based manufacturing of FORCENET provides a foundation for integrating additional modalities, including electrical recording and stimulation^59^, mechanical actuation and cell curvature manipulation^60^, and biochemical sensing^61^, to enable multimodal interrogation of mechanical, electrical, and molecular dynamics in living tissues. By enabling minimally invasive mapping of intercellular force dynamics across tissue-scale, FORCENET provides a framework for studying how mechanical forces are spatially organized, temporally remodeled, and functionally integrated with biochemical signaling during morphogenesis and disease progression.

## Methods

### Device fabrication

The devices were fabricated using SU-8, an epoxy-based negative photoresist. A 50-nm-thick Ni sacrificial layer was first deposited onto a cleaned 4-inch Si wafer by thermal evaporation. To fabricate the force-sensing region, SU-8 2000.5 (Kayaku Advanced Materials) was diluted with cyclopentanone (Sigma-Aldrich) to ∼45% (v/v). The diluted SU-8 was then mixed with one of several fluorescent dyes, including Rhodamine 6G (Sigma-Aldrich), Rhodamine B isothiocyanate (Cayman Chemical), Coumarin 6 (Thermo Fisher Scientific), Rhodamine 110 (Thermo Fisher Scientific), Nile Blue A (Sigma-Aldrich), or Methylene Blue (Sigma-Aldrich), at a final concentration of 100 μg/mL. The mixture was sonicated for 10 min and stored in the dark at room temperature for at least 48 h to ensure complete dye mixing. The fluorescent SU-8 was then spin-coated at 3000 rpm and soft-baked at 65 °C for 1 min and 95 °C for 40 s. The film was exposed to 365-nm ultraviolet (UV) light with a dose of 460 mJ/cm², post-exposure baked at 65 °C for 1 min and 95 °C for 40 s, and developed in SU-8 developer (MicroChem) for 1 min. The patterned structures were subsequently hard-baked at 180 °C for 1 h to enhance crosslinking and mechanical stability. The implantation needle was fabricated in a second lithography step using SU-8 2010 (Kayaku Advanced Materials). SU-8 2010 was spin-coated at 2500 rpm, soft-baked at 65 °C for 1 min and 95 °C for 2 min, exposed to UV light at 320 mJ/cm², post-exposure baked at 65 °C for 1 min and 95 °C for 2 min, and then developed for 3.5 min. Following development, the wafer was hard-baked at 190 °C for 1 h. To release the devices, the wafer was treated with oxygen plasma (100 W, 1 min) and immersed in Ni etchant (Transene) for 15 min to remove the sacrificial Ni layer. Released devices were rinsed three times with deionized water and transferred to 0.1 mg/mL poly-D-lysine solution (Fisher Scientific), where they were stored at 4 °C until use.

### Mechanical characterization

The mechanical properties of SU-8 were characterized by uniaxial tensile testing using a modular force stage (MFS350, Linkam Scientific Instruments) integrated with a small-angle X-ray scattering (SAXS) system (Xenocs Xeuss 3.0). Samples were fabricated from SU-8 2075 (Kayaku Advanced Materials) in a dog-bone geometry (Extended Data Fig. 1a) with a 5-mm-long and 4-mm-wide gauge region and a thickness of approximately 80 μm. During testing, force and displacement were recorded simultaneously to generate stress–strain curves. Engineering stress was calculated from the measured force divided by the cross-sectional area of the gauge region, and engineering strain was calculated from the change in gauge length normalized by the initial gauge length. Young’s modulus was determined from the linear elastic regime of the stress–strain curve using linear regression over the strain range for which the fit exhibited R² > 0.995. To calibrate the mechanical testing system, reference materials with known material properties were measured using the same sample geometry and experimental configuration. A geometric correction factor accounting for systematic offsets in displacement measurements was determined from these reference calibrations and subsequently applied to the SU-8 sample experiments.

### Zebrafish husbandry and transgenic lines

All animal procedures in this study were approved by the Princeton University Institutional Animal Care and Use Committee (IACUC) and performed in accordance with institutional guidelines. Zebrafish (*Danio rerio*) were maintained and bred under standard laboratory conditions. Embryos were maintained at 28.5 °C in E3 medium containing 5 mM NaCl, 0.17 mM KCl, 0.33 mM CaCl_2_, and 0.33 mM MgSO_4_. The following zebrafish lines were used in this study: AB wild-type, Tg(h2az2a:h2az2a-GFP) for ubiquitous nuclear labeling, Tg(actb2:mem-mNeonGreen) for ubiquitous membrane labeling, and Tg(actb2:mem-mNeonGreen; actb2:H2B-mScarlet) for simultaneous membrane and nuclear labeling.

### Device implantation

Following collection, zebrafish embryos were maintained in E3 medium at 28.5 °C and manually dechorionated using fine forceps before reaching the sphere stage. For implantation, embryos were transferred to a glass depression slide containing E3 medium supplemented with methylcellulose to increase viscosity and facilitate manipulation. A 2% (w/v) methylcellulose solution (Sigma-Aldrich) was prepared in E3 medium and kept on ice throughout the experiment. A small drop of methylcellulose was placed onto the glass depression slide and diluted with E3 medium to achieve the desired viscosity. Embryos were transferred onto the methylcellulose using a microcapillary tube (World Precision Instruments) connected to a microinjector and oriented with fine forceps. Individual devices, consisting of a FORCENET sensing region connected to an implantation needle by interconnecting ribbons, were transferred into the same preparation and positioned adjacent to the embryo. Under a stereomicroscope, the implantation needle was guided to a targeted position of the embryonic tissue and advanced through the blastoderm until it traversed the tissue, thereby delivering the FORCENET sensing region into the target location within the embryo. The interconnecting ribbons were then severed using microsurgical scissors and the implantation needle was removed. Following implantation, embryos were incubated in E3 medium for 10 min to dissolve the methylcellulose. Residual methylcellulose was then removed, when necessary, by gently rinsing with E3 medium using a Pasteur pipette. Implanted embryos were subsequently transferred to glass-bottom imaging dishes and maintained at 28.5 °C until imaging.

### Zebrafish imaging

Implanted and non-implanted control zebrafish embryos (4–5 hpf) were mounted in 1% low-melting-point agarose (Sigma-Aldrich) prepared in E3 medium on 35-mm glass-bottom dishes or 12-well glass-bottom plates. Embryos were transferred into liquid agarose using a glass capillary, oriented with forceps before gelation, and immersed in E3 medium throughout imaging. 3D multiphoton imaging was performed using a Nikon AX R multiphoton microscope equipped with a 20× water-immersion objective (NA = 0.95). A tunable femtosecond laser was operated at 960 nm to excite mNeonGreen-labeled membranes and GFP-labeled nuclei, while a 1040-nm laser was used to excite mScarlet-labeled nuclei and Rhodamine B isothiocyanate-labeled FORCENET devices. Fluorescence emission was collected using appropriate bandpass filters. Images were acquired at a lateral sampling size of 0.23–0.74 μm with a z-step size of 2–3 μm. Time-lapse images were collected between 5 and 16 hpf with intervals ranging from 2 to 16 min. Embryos were maintained at 28.5 °C throughout imaging. Image denoising was performed using the Nikon NIS-Elements AI denoising module.

High-resolution imaging of cell mitosis adjacent to FORCENET was performed using a custom built Multimodal Optical Scope with Adaptive Correction (MOSAIC)^40^. Embryos were mounted in 1% agarose/E3 medium at 28.5 °C and positioned to allow optical access to the implanted device region. 3D time-lapse imaging was performed between 6 and 8 hpf with time intervals approximately 10 s to capture cell division events adjacent to the device. Fluorescence excitation was performed at 488 nm for nuclear GFP and 560 nm for Rhodamine B isothiocyanate-labeled FORCENET. Volumetric images were acquired with a lateral sampling size of 108 nm and a z-step size of 200 nm. Images acquired in sample stage scanning mode were deskewed and deconvolved using the PetaKit5D package^62^.

Whole-embryo developmental imaging was performed using a Zeiss Axio Observer Z1 inverted fluorescence microscope equipped with a 5× air objective. Correlative brightfield and fluorescence images were acquired every 10 min and overlaid for visualization. Embryos were maintained at 28.5 °C throughout imaging.

### Analysis of embryo survival and morphology

To assess whether FORCENET implantation and integration perturb embryonic development, morphological parameters were quantified in implanted and non-implanted wildtype (WT) control embryos at sphere and shield stages. Embryos were imaged using a Zeiss SteREO Discovery.V8 stereomicroscope, and anatomical landmarks were manually annotated in Fiji^63^.

Blastoderm morphology was characterized by measuring the central blastoderm height (H), defined as the distance between the yolk surface and blastoderm margin at the embryo midline, together with corresponding left (H_L_) and right (H_R_) heights measured at positions one-quarter of the embryo width from the midline. Blastoderm asymmetry was quantified as:

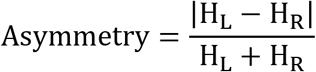

This provides a dimensionless metric independent of embryo size. Statistical comparison of implanted and control embryos was performed using an unpaired t-test for blastoderm height and a two-sided Mann–Whitney U test for asymmetry.

To evaluate embryo viability, implanted and control embryos were kept at 28.5 °C under standard conditions until 24 hpf and were classified as developed normally, developed abnormally, or did not survive based on stereomicroscopic examination.

### Comparison of cell density and kinematics near and away from FORCENET

To quantitatively determine whether FORCENET locally perturbs tissue organization, cellular properties adjacent to the device were compared with anatomically matched contralateral regions. Device-adjacent regions (+FORCENET) were defined by dilating the segmented FORCENET structure by 20–30 μm from its centerline and masking the resulting volume using a cellular foreground segmentation in Labkit^64^. Contralateral regions (−FORCENET) were generated by reflecting the device segmentation across the embryo’s anatomical plane of symmetry, identified from the 3D multiphoton images at 10–12 hpf. This approach produced spatially matched regions that experienced comparable developmental trajectories while differing only in the presence or absence of the implanted device.

### Extraction of device deformation from volumetric images

We developed a custom computer-vision pipeline to extract nodal positions and rotations from 3D multiphoton image stacks (Supplementary Video 4). First, FORCENET was segmented using Labkit^64^ and skeletonized to obtain the centerlines of the force-sensing beam elements. Beam intersections corresponding to nodes were identified from the local connectivity of the skeleton. Candidate node positions were initially localized at voxel resolution and subsequently refined using fluorescence intensity profiles extracted along the axial direction. To reduce localization errors arising from anisotropic voxel sampling, Gaussian mixture models were fitted to the intensity distributions, enabling sub-voxel estimation of nodal coordinates and resolution of overlapping structural features. All reconstructed node positions were manually verified using BigWarp in Fiji.

Next, we extracted internodal positions for calculation of nodal rotations. The centerline of each beam element was reconstructed by sampling a series of cross-sectional planes along the beam connecting neighboring nodes. Within each plane, candidate beam center positions were identified by fitting one- or two-component Gaussian mixture models to the fluorescence intensity distribution. Because multiple candidate trajectories can arise in densely packed regions, the optimal beam centerline was determined using a Viterbi-based path optimization that penalized discontinuous trajectories and excessive curvature. The midpoint along the reconstructed beam centerline was defined as an internodal landmark.

The reconstructed nodal and internodal coordinates were then registered to the undeformed device geometry using rigid-body alignment, which minimized the mean-squared displacement between corresponding points while accounting for global translations and rotations of the device. Nodal displacements were obtained directly from the registered coordinates. To estimate nodal rotations, the measured nodal and internodal positions were imposed as displacement constraints in a finite element model of the undeformed lattice, with rotational degrees of freedom left unconstrained. The resulting finite element solution provided the nodal rotations used for subsequent force reconstruction.

### Finite element modeling and stress reconstruction

FORCENET was modeled as a network of elastic beam elements connected by nodes corresponding to the honeycomb lattice geometry. The analysis assumed mechanical equilibrium during imaging. Because the beam thickness (∼180 nm) is substantially smaller than its width (∼3 μm), deformations were assumed to be dominated by bending along the beam thickness direction. Stress reconstruction was performed using an FEA framework based on PyNite. Specifically, measured nodal translations and rotations were related to nodal forces through the global stiffness matrix (Extended Data Fig. 6a):

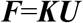

where ***F*** is a 6*N* × 1 vector, with *N* denoting the number of nodes; each node contributes three force components and three moment components. ***U*** is a 6*N* × 1 vector containing the experimentally measured nodal displacements and rotations, with three displacement components and three rotational components for each node. ***K*** is the structural stiffness matrix with dimensions 6*N* × 6*N*. To mitigate spatial registration errors, we swept beam thickness parametrically, fit the reconstructed stress estimates as a polynomial function of thickness, and retained only the cubic term expected for rectangular beam geometries. The reconstructed nodal forces were subsequently converted into effective distributed loads on individual beam elements by assigning that the force at each node was shared equally among the beam elements connected to that node. For the beam element connecting nodes *i* and *j*, the distributed load was estimated as:

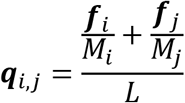

where ***q***_i,j_ is the distributed load vector, ***f***_i_ and ***f***_j_ are the three-component nodal force vectors at node *i* and *j, M*_i_ and *M*_j_ are the numbers of beams elements connected to these nodes, and *L* is the beam length. The effective stress along the beam thickness direction was then estimated as:

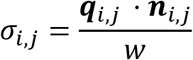

where ***n***_i,j_ is the unit normal vector along the beam thickness direction and *w* is the beam width. This conversion reports the effective stress averaged over the beam element. Portions of the device extending into the agarose support were modeled as fully fixed boundaries, with all translational and rotational degrees of freedom constrained.

### Intercellular stress interpolation

To convert the discrete element stress map into a continuous intercellular stress field within the FORCENET sensing region, we performed piecewise-linear barycentric interpolation over a triangulated representation of the honeycomb lattice. First, the reconstructed beam element stresses were mapped to the vertices of the honeycomb lattice by averaging the stress values of all beam elements connected to each vertex. To avoid large or poorly conditioned triangles during interpolation, an additional interpolation node was placed at the centroid of each honeycomb cell. The stress value at this centroid node was defined as the mean stress of the six surrounding honeycomb vertices. Together, the six vertices and the centroid nodes partitioned each honeycomb cell into six triangles. Additional boundary triangles were constructed along the FORCENET perimeter to complete the triangulation grid. The continuous stress field was then evaluated at any point **r** located inside a triangle **T**_ijk_ = (***r***_i_, ***r***_j_, ***r***_k_) using linear interpolation in barycentric coordinates:

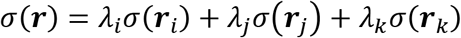

where *λ*_i_, *λ*_j_, *λ*_k_ are the barycentric weights of ***r*** with respect to the trigangle **T**_ijk_, satisfying:

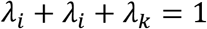

and

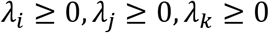

Points outside the triangulated FORCENET sensing region were excluded. The interpolated stress field was then resampled onto a regular grid with 1 μm spacing for visualization and downstream analysis, including autocorrelation and cross-correlation characterizations.

### Spatial autocorrelation analysis

To quantify the spatial organization of intercellular stresses, reconstructed stress fields were projected onto the undeformed device geometry and interpolated using barycentric interpolation. Spatial autocorrelation functions were computed from mean-subtracted stress fields and corrected for finite-image overlap at increasing spatial lags. Directional correlation lengths were determined from radial profiles extracted from the autocorrelation map. The correlation length (*ξ*) was defined as the first zero-crossing of the autocorrelation function. Profiles that did not cross zero within the analyzed domain were excluded. Correlation lengths were calculated for 50 directions spanning −90° to 90°, with −90°/90° corresponding to lateral/medial and 0°/180° corresponding to anterior/posterior directions of the embryos. Maximum, minimum, median, and mean correlation lengths, together with the orientation and ellipticity of the autocorrelation field, were subsequently quantified.

### Cell behaviors quantification and correlation with intercellular stresses

To quantify cell density, speed, and acceleration magnitude, nuclei were segmented from 3D image stacks using Cellpose-SAM^43^ and tracked over time using Ultrack^44^. Cell density was calculated as the number of segmented nuclei within a given volume. Cell positions were defined as nuclear centroids. Velocities and accelerations were calculated from finite differences of cell positions and subsequently interpolated onto the original imaging time points. Speed and acceleration magnitude were defined as the magnitudes of the corresponding vectors.

To correlate cell behaviors with intercellular stresses, stresses, cell density, velocity, and acceleration were converted into scalar fields for spatial correlation analysis. Stress values were assigned to beam internodes and interpolated onto a regular 5 × 5 × 5 μm^3^ grid using radial-basis-function interpolation using a linear kernel:

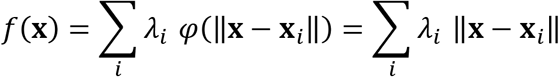

where {**x**_*i*_} are internodes positions and the weights {*λ*_*i*_} are obtained by solving the interpolation system exactly without no smoothing. In this way, the interpolated scalar field reproduces the sampled values at the corresponding internodes. Cellular behaviors (density, speed, and acceleration magnitude) were computed within 40 × 40 × 40 μm^3^ voxels and subsequently interpolated onto the same grid using the same radial-basis-function interpolation. Spatial cross-correlations were calculated between mean-subtracted fields after masking regions within 20–30 μm of the device. Cross-correlation functions were corrected for the decreasing overlap volume at increasing spatial lags and normalized by the standard deviations of the corresponding variables.

### Pharmacological inhibition of FGF signaling

FGF signaling was inhibited using SU5402, a small-molecule inhibitor of FGF receptor tyrosine kinases. A 10 mM stock solution of SU5402 in dimethyl sulfoxide (DMSO) was obtained from Sigma-Aldrich and diluted into E3 medium to a final concentration of 10 μM immediately before use. Dechorionated embryos were treated with the SU5402-containing medium beginning at 30% epiboly and maintained in the medium throughout imaging. Control embryos were not treated and maintained in E3 medium throughout imaging. All treatments were performed under reduced-light conditions to minimize photodegradation of the compound.

### Cerebral organoid generation

Human induced pluripotent stem cells (hiPSCs) were maintained under feeder-free conditions in mTeSR™ Plus medium (STEMCELL Technologies) on vitronectin-XF–coated plates and passaged every 4–6 days using 0.5 mM EDTA (Thermo Fisher Scientific). Following passaging, cells were cultured in medium supplemented with 10 μM Y-27632 (ROCK inhibitor; STEMCELL Technologies, Canada) for the first 24 h to enhance cell survival. Human cerebral organoids were generated from hiPSCs following established protocols^65^ with minor modifications. Briefly, hiPSCs were dissociated with Accutase (Sigma) and seeded at 9,000 cells per well in low-attachment 96-well U-bottom plates in mTeSR™ Plus medium containing 20 μM Y-27632 to form embryoid bodies. Neural induction was performed from days 1–5 in Essential 6™ medium (Gibco) supplemented with 5 μM dorsomorphin (Sigma) and 10 μM SB-431542 (Tocris). On day 6, spheroids were transferred to neural medium consisting of Neurobasal-A (Gibco) supplemented with B27 minus vitamin A (Invitrogen), GlutaMAX (Invitrogen), and penicillin-streptomycin (Sigma), together with 20 ng/mL bFGF (Peprotech) and 20 ng/mL EGF (Peprotech). Organoids were cultured on an orbital shaker starting on day 18 at 60–80 rpm. Beginning on day 25, organoids were transferred to low-attachment 24-well plates and maintained in neural medium supplemented with 20 ng/ml BDNF (Peprotech) and 20 ng/mL NT-3 (Peprotech). Medium was refreshed daily or every other day throughout culture.

## Acknowledgements

We thank S. Megason and A. Murphy for the membrane and two-color membrane-nuclei fish lines, N. Claussen, D. Cohen, M. Haataja, A. Košmrlj, B. Reneker, C. Warren for helpful discussions, C. Brangwynne, G. Laevsky and S. Wang for help with the imaging, and the Laboratory Animal Resources at Princeton University for their help in zebrafish maintenance. This work was performed in part at the Micro/Nano Fabrication Center (MNFC) and the Imaging and Analysis Center (IAC) at the Princeton Materials Institute (PMI). T.-M. F. acknowledges the support from National Science Foundation CAREER award (2543120), National Institutes of Health MIRA award (R35GM165761), and Princeton School of Engineering and Applied Science Innovation fund; Y.L. and T-M.F. acknowledge the support from National Institute of Health grant R21AG095506; R.D.B. acknowledges the support from New Jersey Commission on Cancer Research; S.K. acknowledges the support from Gilbert S. Omenn, M.D., Ph.D. ’61 & Martha Darling *70 Fellows; X.Z. acknowledges the support from Omenn-Darling Bioengineering Institute Innovators (ODBI²) Fellowship.

## Author contributions

T.-M.F conceived the idea and supervised the project. F.L., S.K., and X.Z. performed the device fabrication and implantation. F.L., S.K., V.G., C.B.Y., and R.D.B. prepared the zebrafish. J.W. and Y.L. prepared the organoids. F.L., S.K., and T.-M.F. performed the live fluorescent imaging. J.D.W., Y.Q., M.J.L. developed the imaging processing pipeline. F.L., S.K., G.M., and A.Y. collected the anatomical data and performed the analysis. J.D.W. and F.L. visualized the data. J.D.W. performed the cell segmentation and tracking. J.D.W., S.K., Z.S., and H.A.S. performed the mechanical modeling. F.L., J.D.W., and T.-M.F. prepared the figures and wrote the manuscript with input from all authors.

## Data and materials availability

All data needed to evaluate the conclusions are present in the paper and Supplementary Materials.

## Supplementary Information

Supplementary Notes 1-6

Supplementary Figure 1

Legends for Supplementary Videos 1-7

Supplementary References

## Extended Data Figures and Figure Legends

**Extended Data Fig. 1.**
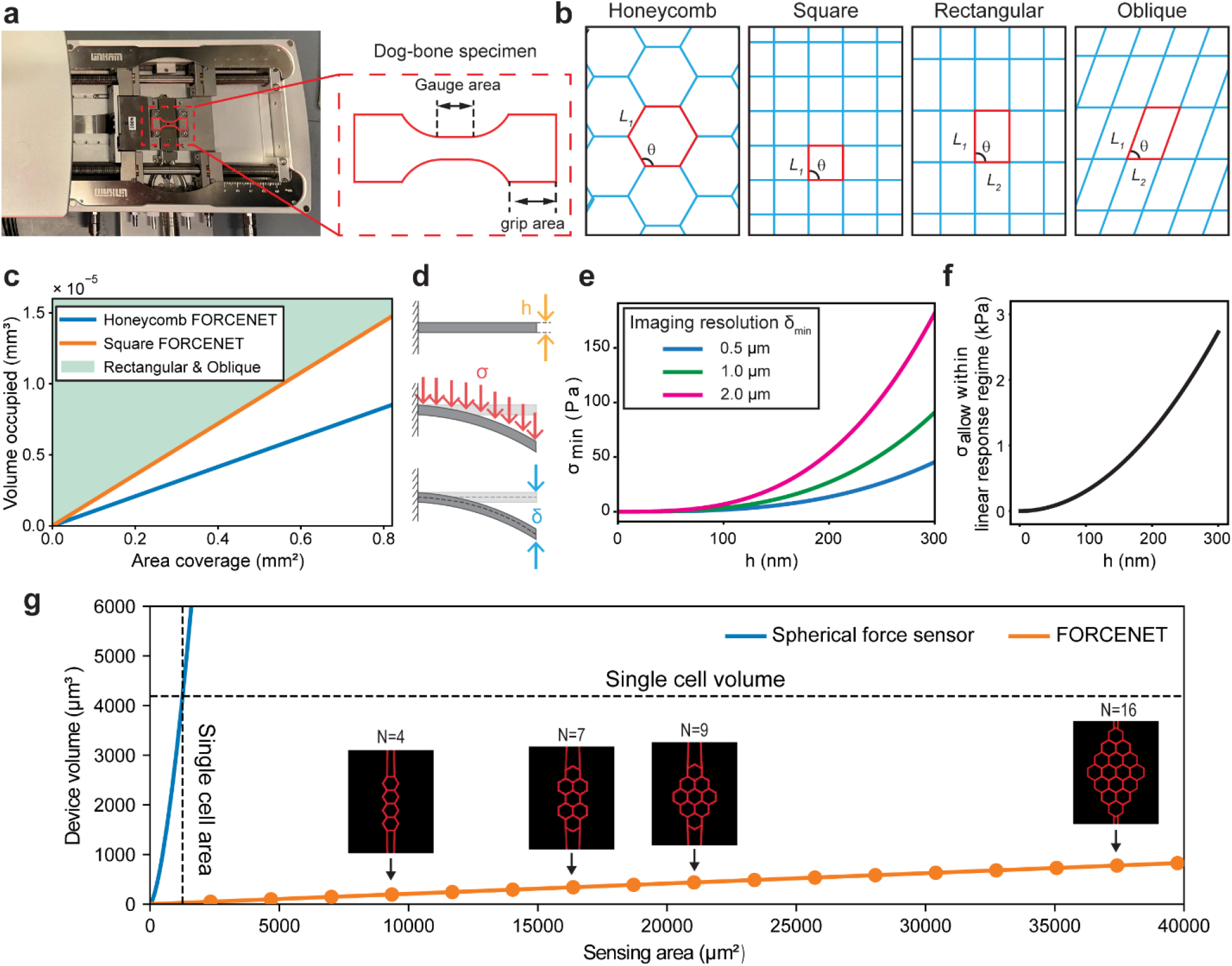
Design principles and mechanical optimization of FORCENET. **a,** Experimental setup for tensile testing of SU-8 specimens. Left, photograph of the tensile testing stage with a schematic sample outline overlaid. Right, schematic of the dog-bone specimen geometry, indicating the gauge area used for strain measurement and the grip area used for mounting. **b,** Schematic of four candidate lattice structures for FORCENET design: honeycomb, square, rectangular, and oblique. Blue lines indicate the lattice network, and red outlines highlight representative unit cells. Geometric parameters used to define the unit-cell dimensions and orientation are indicated. Labels *L*_1_ and *L*_2_ denote the lattice dimensions, and *θ* denotes the lattice angle. **c,** Plots demonstrating how tissue volume occupied by FORCENET scales with sensing area covered by FORCENET for different FORCENET lattice structures. **d,** Schematic of the mechanical model for each FORCENET element. Each element is modeled as a cantilever beam with one end fixed and subjected to uniformly distributed loading. Here, *h* denotes the beam thickness, σ the applied uniform stress, and δ the maximum deflection. **e,** Minimum detectable stress (σ_min_) as a function of beam thickness for different imaging resolutions, calculated from the cantilever model. **f,** Maximum measurable stress (σ_allow_) within the linear elastic regime of SU-8 defined in Fig. 1c, plotted as a function of beam thickness based on the cantilever model. **g,** Scaling of device volume with sensing area for spherical force sensors and FORCENET. Insets show representative FORCENET designs with different numbers of honeycomb unit cells. The vertical dashed line indicates the estimated single-cell surface area, and the horizontal dashed line indicates the estimated single-cell volume.

**Extended Data Fig. 2.**
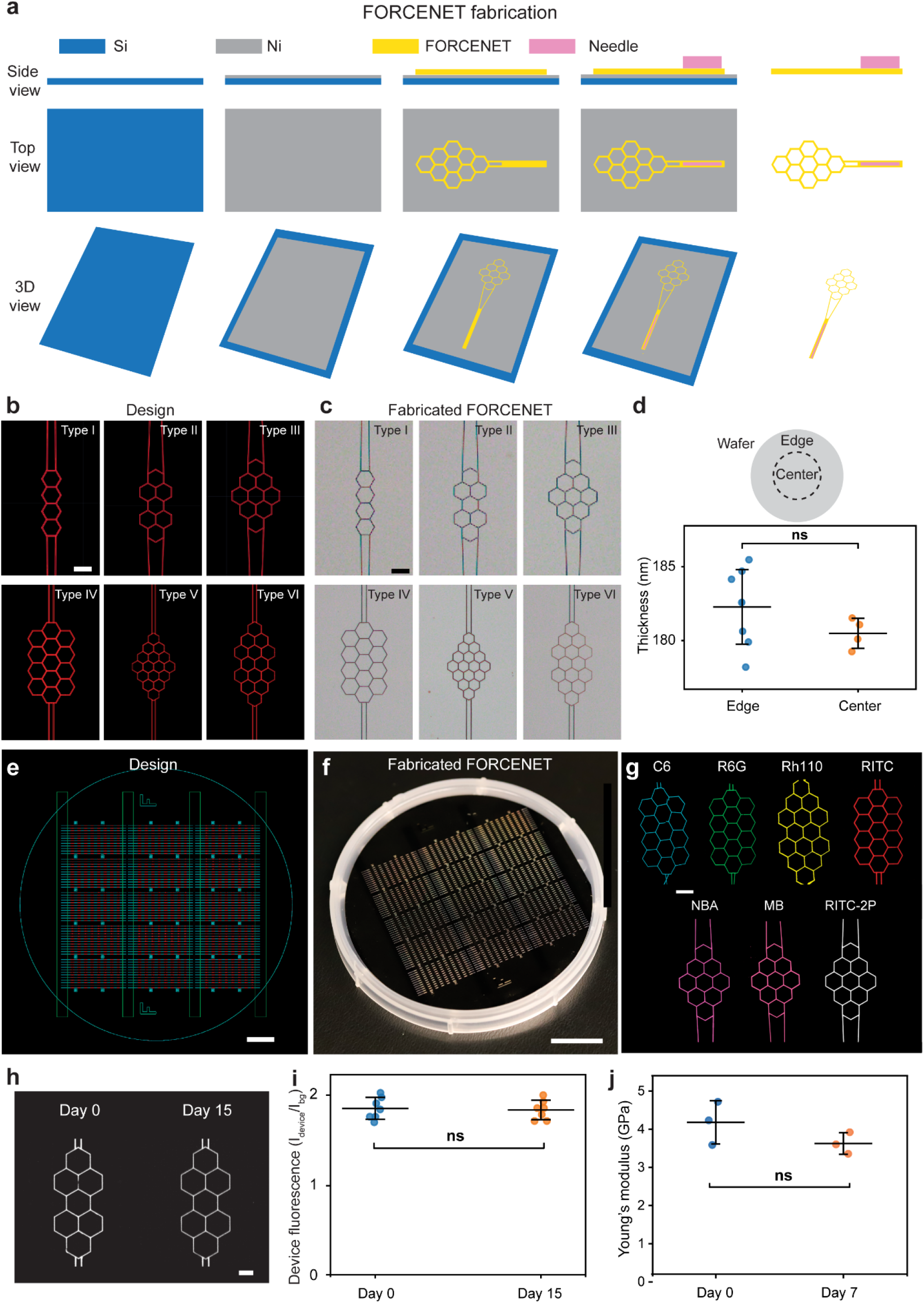
Fabrication, optical labeling, and stability characterization of FORCENET. **a,** Schematic illustration of the fabrication process for FORCENET. Side views (top panels), top views (middle panels), and 3D views (bottom panels) show sequential steps, including deposition of a Ni sacrificial layer (gray) on a Si substrate (blue), patterning of the FORCENET structure (yellow), fabrication of the penetrating needle (pink), and release of the device by etching the Ni layer. **b,** CAD layouts of structures with different beam sizes and honeycomb-cell arrangements. **c,** Corresponding bright-field images of the fabricated devices. Scale bars, 50 μm. **d,** Thickness uniformity across the wafer. Thickness was measured at edge and center regions, as illustrated in the schematic. Data are presented as mean ± s.d.. Each point represents an individual measurement. No significant difference (ns) was observed between groups (two-tailed unpaired t-test, p = 0.113). **e,** Three-layer photomask layout showing the arrangement of >2,000 devices across a 4-inch wafer. Scale bar, 10 mm. **f,** Photograph of the fabricated wafer containing arrays of FORCENET. Scale bar, 20 mm. **g,** Representative fluorescence images of FORCENET labeled with different dyes (C6, Coumarin 6; R6G, Rhodamine 6G; Rh110, Rhodamine 110; RITC, Rhodamine B isothiocyanate; NBA, Nile Blue A; MB, Methylene Blue; RITC-2P, RITC imaged with two-photon microscopy). All images besides RITC-2P were captured using confocal microscopy. Scale bar, 50 μm. **h,** Representative fluorescence images showing FORCENET at day 0 and day 15 after fabrication. Scale bar, 30 μm. **i,** Normalized device fluorescence intensity measured at day 0 and day 15, where *I*_device_ is the intensity of the device and *I*_bg_ is the intensity of the background. Each dot represents one device; bars indicate mean ± s.d. No significant difference was observed between groups (two-tailed Welch’s t-test, p = 0.786). **j,** Young’s modulus of SU-8 measured at day 0 and after 7 days of immersion in E3 medium at 28.5 °C. Values were calculated from the linear region of the stress–strain curve, defined by a linear fit with R^2^ > 0.995. Points represent individual samples; bars indicate mean ± s.d. Statistical significance was assessed using a two-tailed Welch’s t-test (ns, not significant, p = 0.230).

**Extended Data Fig. 3.**
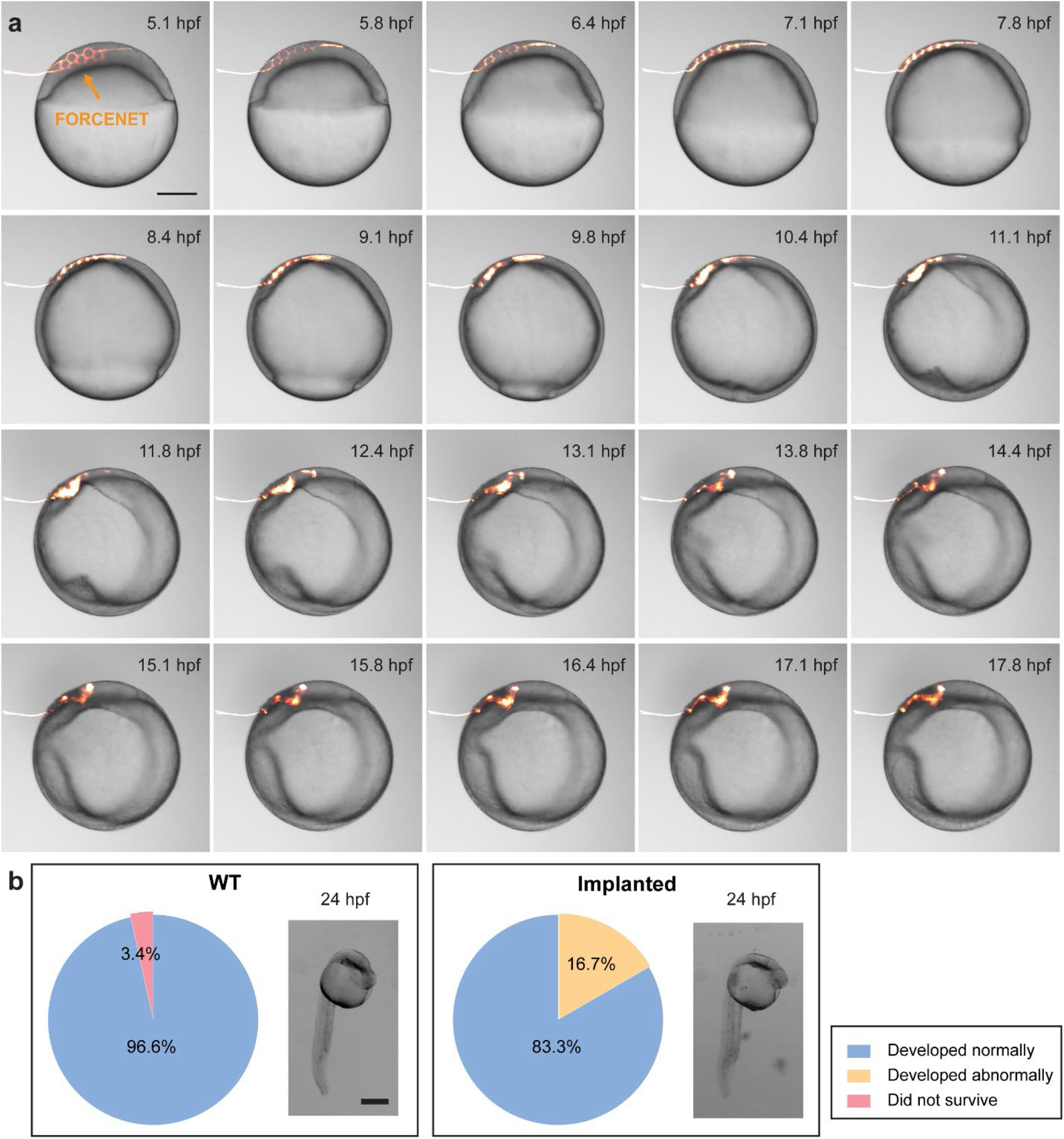
FORCENET integrates into embryos without disrupting normal development. **a,** Correlative brightfield-fluorescence images (see also Supplementary Video 1) showing the development of an embryo with implanted FORCENET (orange). Scale bar, 200 μm. **b,** Survival rate of non-implanted WT control (n = 29) and implanted embryos (n = 12) at 24 hpf. Pie charts show the percentages of embryos that developed normally (blue), abnormally (orange), and did not survive (pink). Insets show representative brightfield images at 24 hpf. Scale bar, 300 μm.

**Extended Data Fig. 4.**
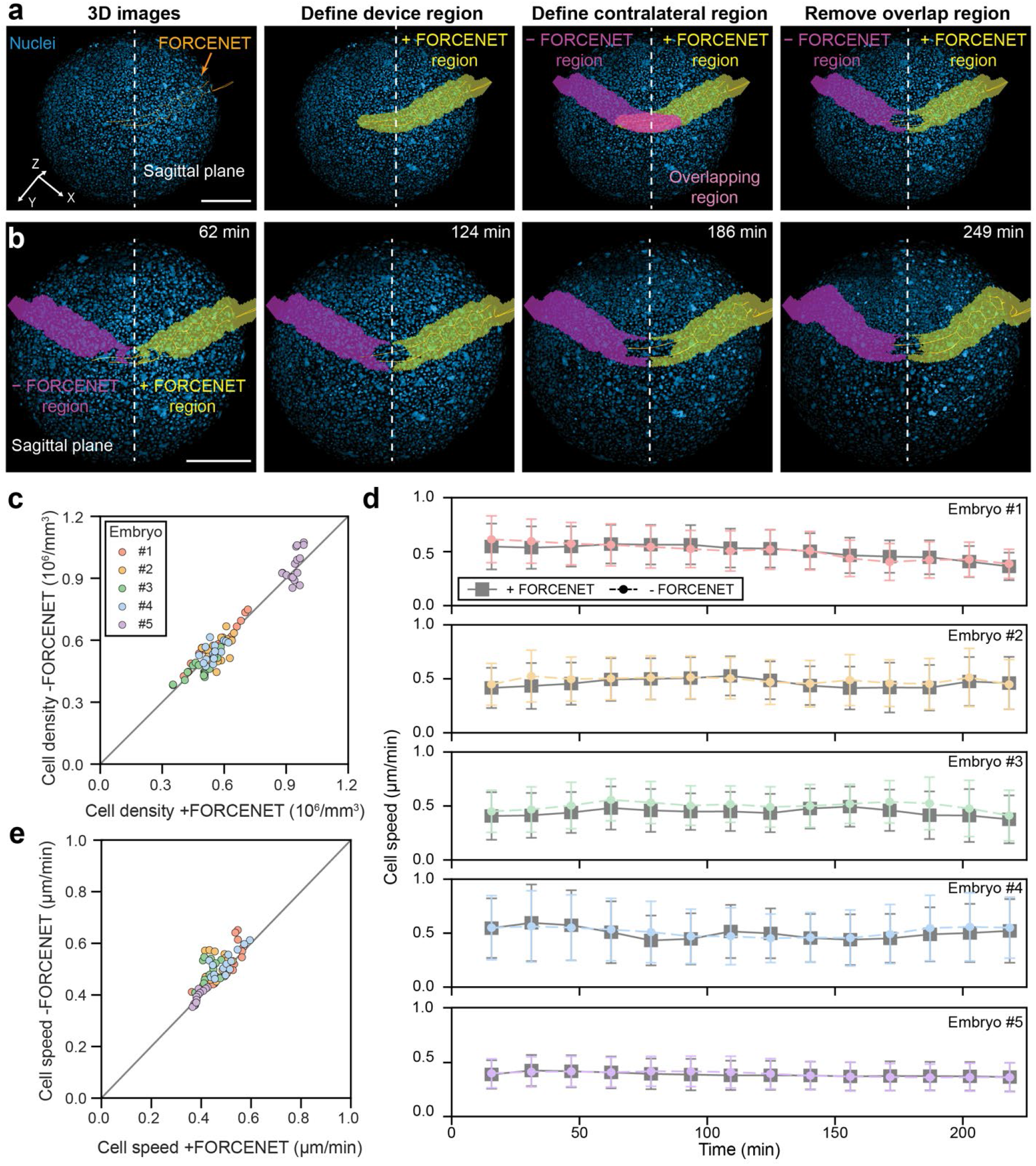
FORCENET does not measurably alter local cell density or cell speed. **a,** Workflow for defining FORCENET-implanted (+FORCENET) and contralateral (−FORCENET) regions across an anatomical plane. Raw 3D imaging data show nuclei (blue) and FORCENET (orange), with the sagittal plane indicated by white dashed line. The device region is segmented (yellow), mirrored across the sagittal plane to define a contralateral region (magenta), and overlapping regions are removed to obtain paired regions for analysis. Scale bar, 200 μm. **b,** Time-lapse visualization of the temporal evolution of +FORCENET (yellow) and −FORCENET (magenta) regions in a representative embryo. Scale bar, 200 μm. **c,** Comparison of cell density in +FORCENET and −FORCENET regions across five embryos. Each dot represents one time point, color-coded by embryo. The gray diagonal indicates equality between the two regions. **d,** Time-resolved cell speed measured in +FORCENET and −FORCENET regions for each embryo. Solid gray lines with square markers indicate the mean cell speed in +FORCENET regions, and dashed colored lines with circular markers indicate the mean cell speed in −FORCENET regions. Error bars represent ± s.d.. **e,** Comparison of cell speed measured in +FORCENET and −FORCENET regions across five embryos. Each dot represents one time point, color-coded by embryo. The gray diagonal indicates equality between the two regions.

**Extended Data Fig. 5.**
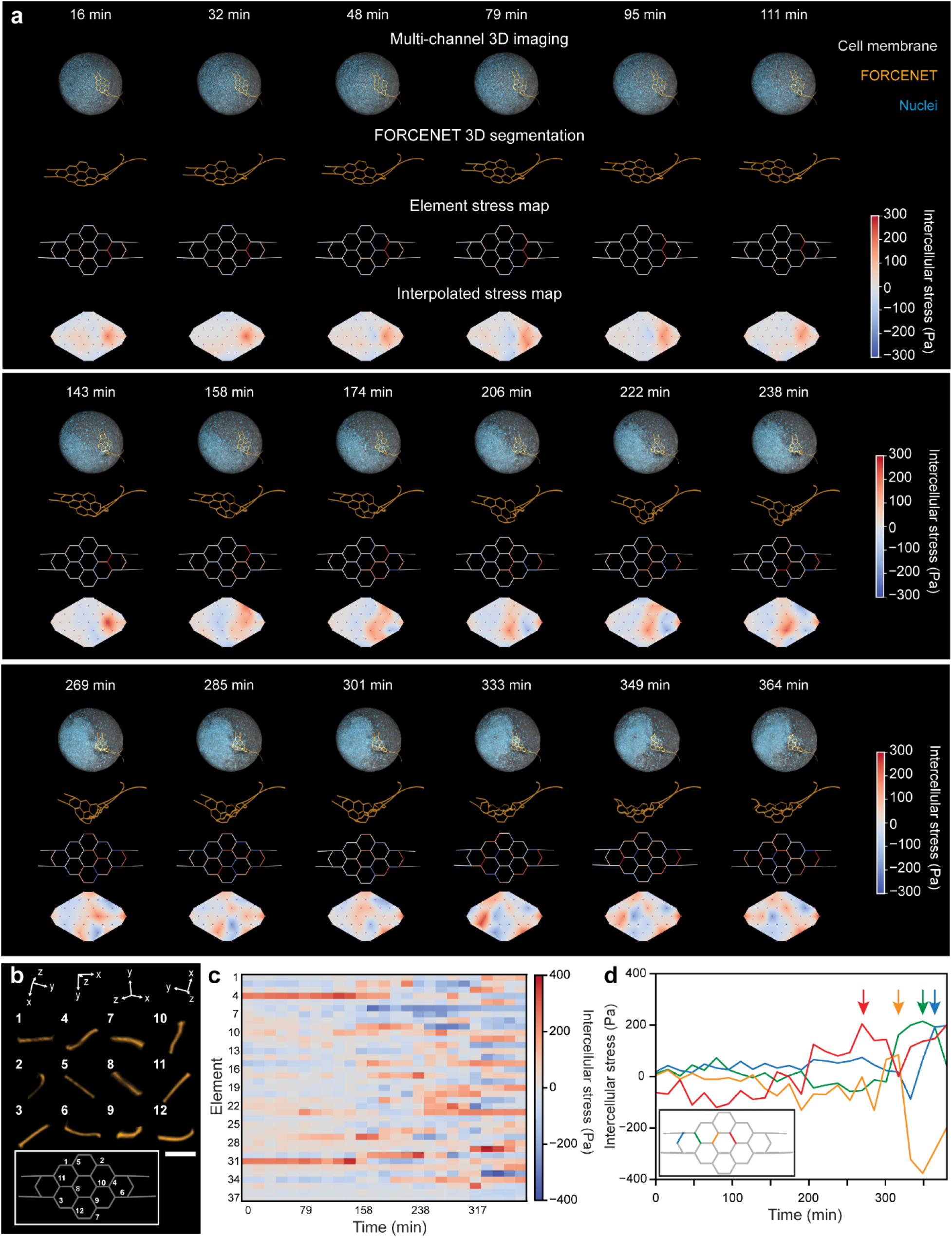
Dynamic intercellular stress mapping during zebrafish gastrulation. **a,** Multi-channel 3D images of an embryo with implanted FORCENET, corresponding FORCENET 3D segmentations, intercellular stress mapped onto beam elements, and interpolated spatial distributions of intercellular stress across time points, shown from top to bottom. Colors in the multi-channel imaging correspond to nuclei (blue), cell membrane (gray), and FORCENET (orange). **b,** Representative segmented beam elements showing heterogeneous deformation across the force-sensing network. Inset is a schematic of the corresponding beams on FORCENET. Scale bar, 25 μm. **c,** Heatmap representation of intercellular stress for individual FORCENET elements over time. Each row corresponds to a beam element and each column to a time point. Color bar indicates intercellular stress. **d,** Temporal evolution of intercellular stress for representative elements. Arrows indicate the maximum intercellular stress for each element. Inset depicts the elements corresponding to the curves.

**Extended Data Fig. 6.**
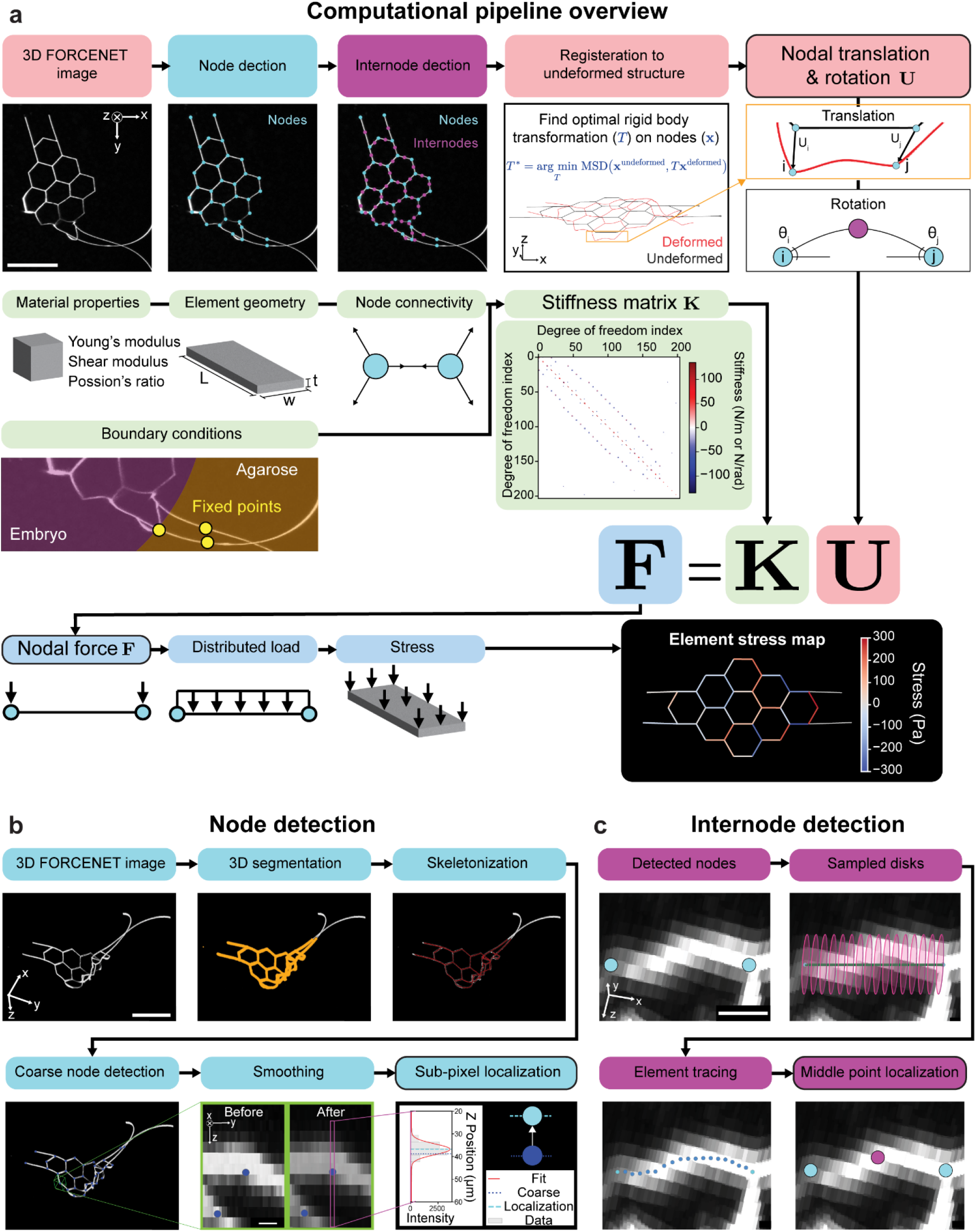
Image-processing and computational pipeline for FORCENET stress reconstruction. **a,** Computational pipeline overview. 3D images of FORCENET were used to extract node and internode coordinates, which were registered to the undeformed configuration to determine nodal displacements and rotations. A finite-element stiffness matrix was then constructed from the measured material properties, device geometry, connectivity, and boundary conditions. Nodal forces were computed from the stiffness matrix and the nodal displacements and rotations, then converted into distributed loads and intercellular stresses to generate an element stress map. **b,** Node detection pipeline. 3D FORCENET images were processed through segmentation and skeletonization, followed by coarse node detection. Gaussian smoothing and sub-pixel localization are applied to refine node positions. **c,** Internode detection pipeline. For each node pair, a series of disks was sampled along the straight line connecting the nodes. Candidate beam center positions were identified within each disk by Gaussian fitting the cross-sectional fluorescence intensity profile. These candidate points are then connected using a shortest-path optimization to reconstruct the beam trajectory, and the midpoint along the resulting path was defined as the internode location.

**Extended Data Fig. 7.**
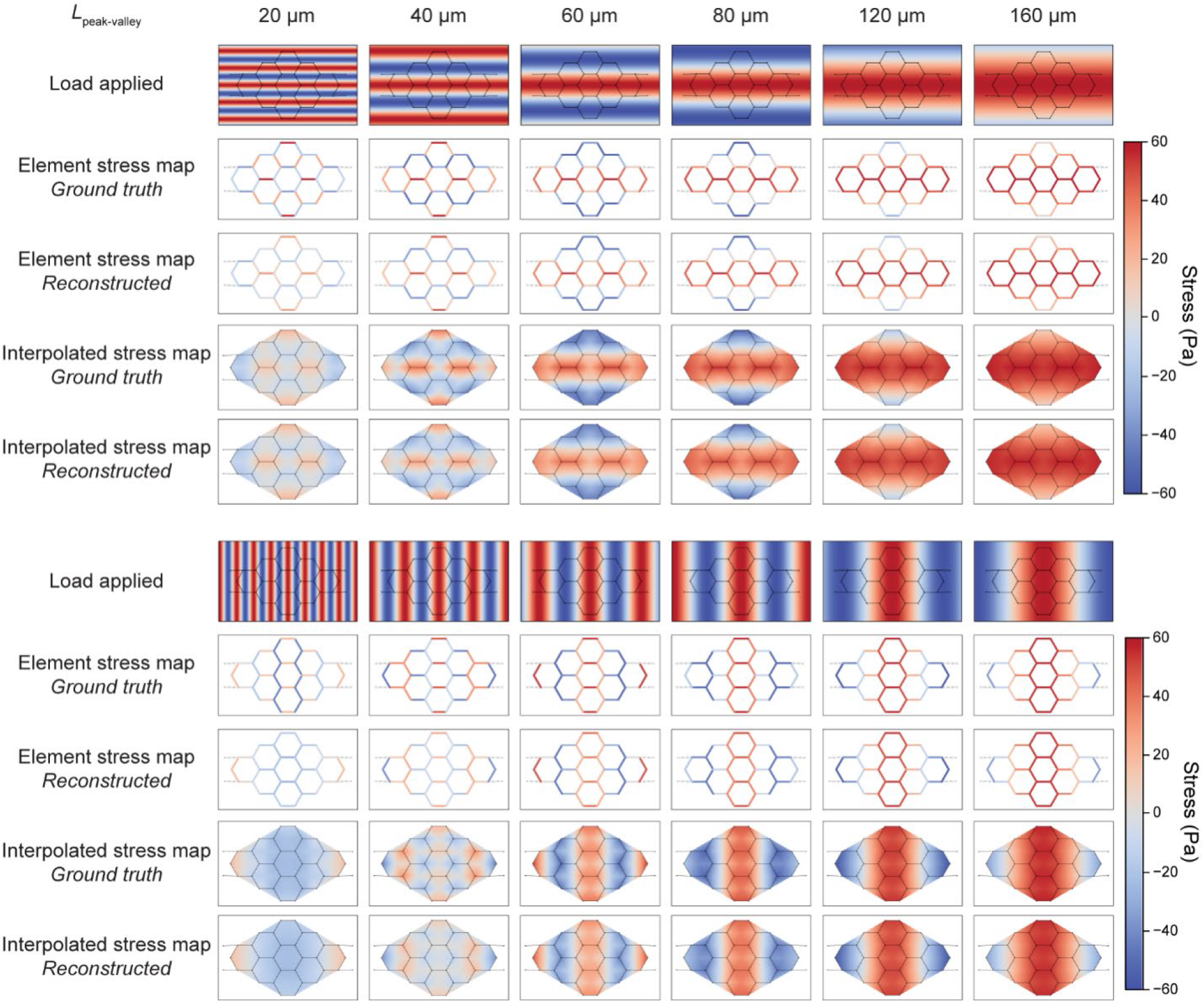
Characterization of FORCENET stress reconstruction across spatial frequencies and orientations. Grating stress fields with different peak-to-valley spacings, *L*_peak-valley_, and two orthogonal orientations were applied to a FORCENET. Forward finite-element analysis was used to obtain the resulting FORCENET deformation, and the nodal translations and rotations from the simulation were used as inputs for the reconstruction pipeline. For each applied stress field, the ground-truth element stress map was obtained by averaging the applied stress over each force-sensing beam element. Element-wise stress maps and spatially interpolated stress maps are shown for both the ground truth and reconstructed stresses.

**Extended Data Fig. 8.**
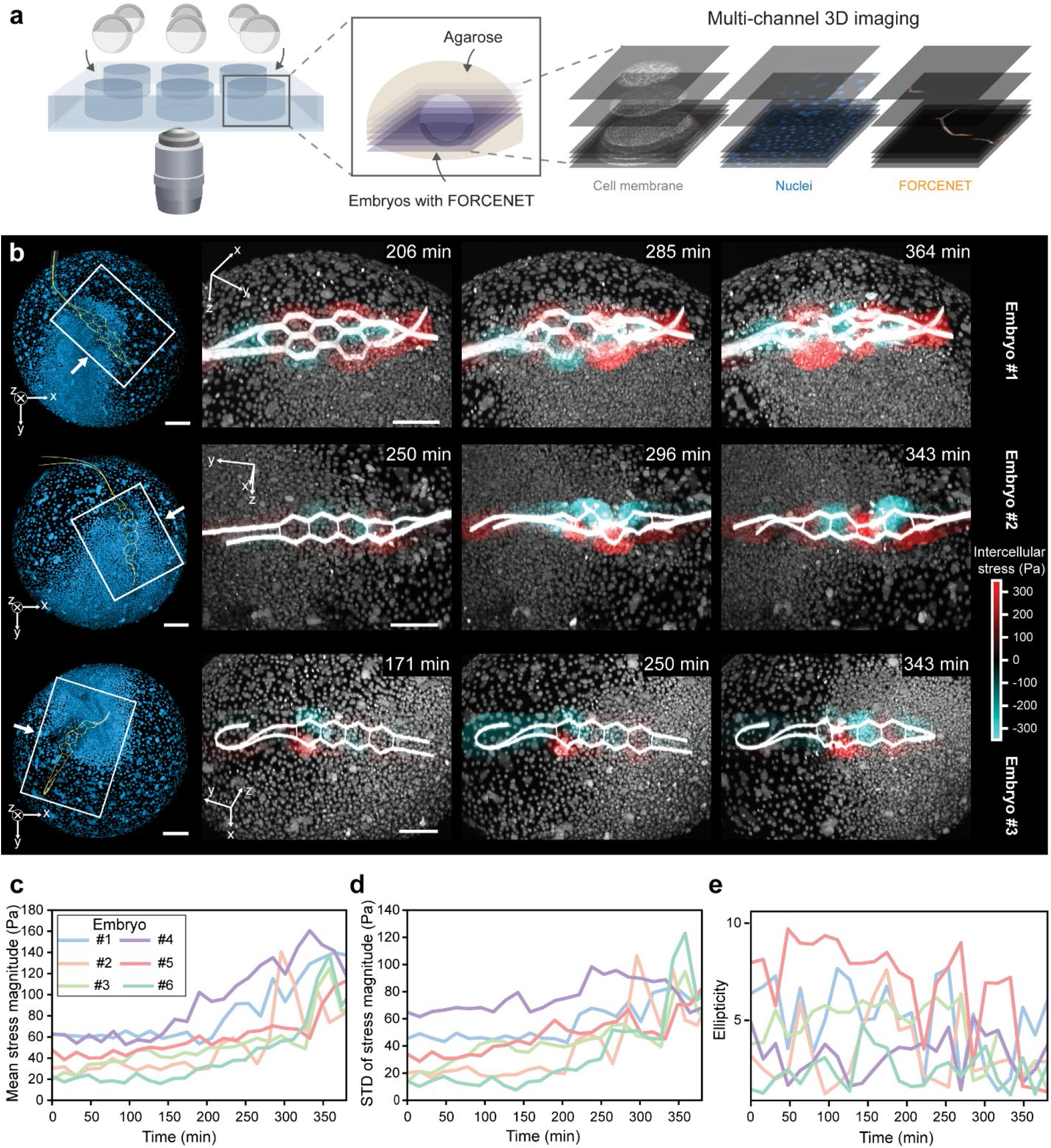
Intercellular stress dynamics across multiple embryos during zebrafish gastrulation. **a,** Workflow for multi-embryo *in vivo* imaging. Multi-channel 3D two-photon imaging captures cell membranes, nuclei, and FORCENET during development. **b,** Time-lapse images and interpolated intercellular stress fields of three representative embryos (rows), each with an implanted FORCENET. Left column shows 3D whole-embryo rendering with nuclei (blue) and FORCENET (orange); remaining columns show time-lapse stress maps zoomed in on the local behavior near FORCENET and nuclei, both in white. Panels **c**–**e** contain analysis performed across a cohort of 6 embryos, with each color representing data from different embryos. **c,** Mean intercellular stress magnitude across all embryos over time. **d,** Standard deviation of the intercellular stress magnitude across all embryos over time. **e,** Temporal evolution of the ellipticity of the spatial autocorrelation field across all embryos.

**Extended Data Fig. 9.**
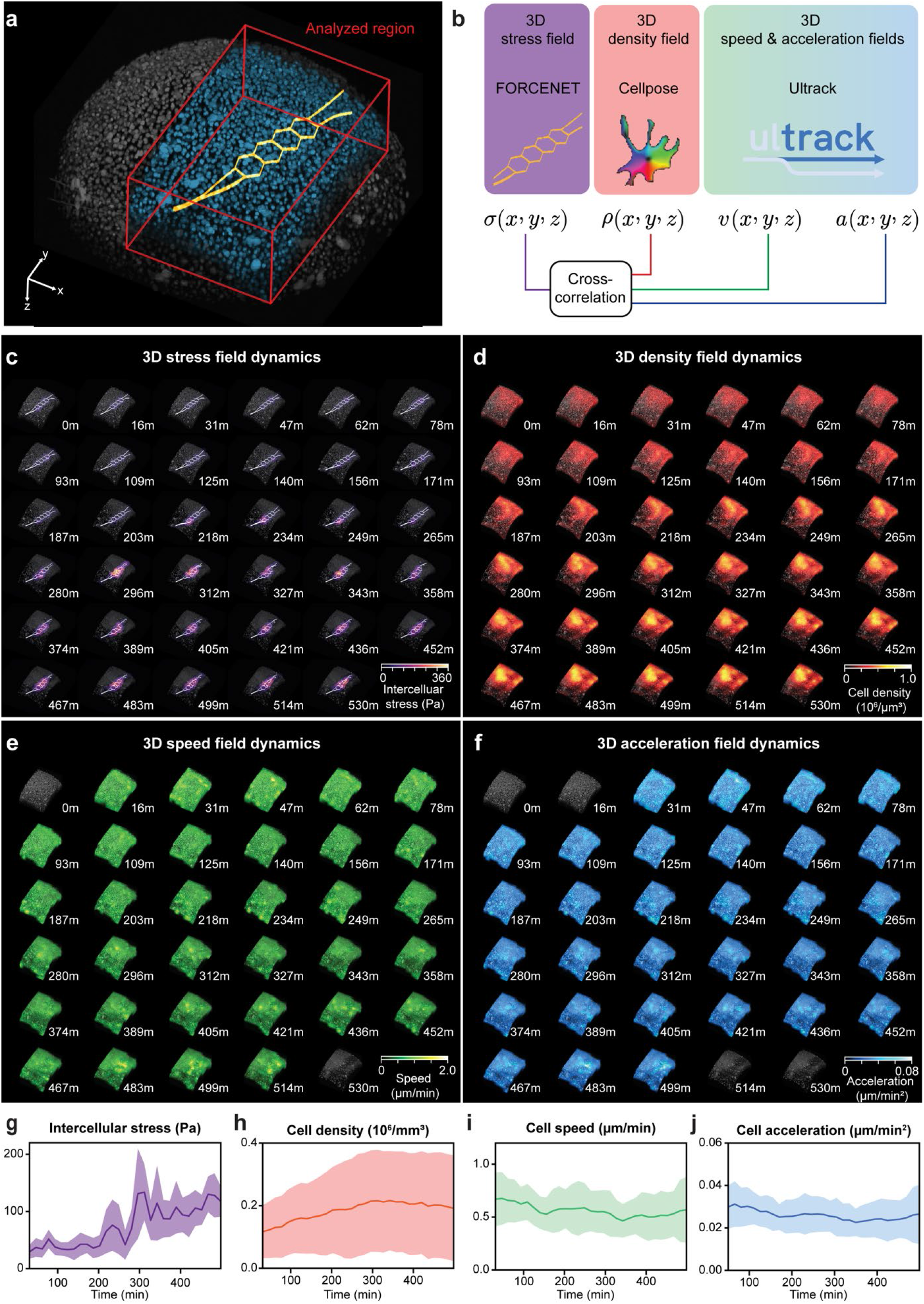
Simultaneous mapping of intercellular stress and cellular dynamics during zebrafish gastrulation. **a,** A sub-volume containing FORCENET (orange) is selected within the embryonic blastoderm. Bounding box dimensions: 75 × 427 × 190 µm^3^ (z × y × x). **b,** Schematic of cross-correlation analysis pipeline. 3D intercellular stress field (*σ*) was measured by FORCENET, cell density field (*ρ*) was obtained from cell segmentation using Cellpose-SAM^43^, velocity field (*v*) and acceleration field (*a*) were computed from cell tracking using Ultrack^44^. Cross-correlation analyses were then performed to quantify the relationship between intercellular stress and cell density and cellular dynamics. (**c**–**f**) Time-lapse visualization of spatiotemporal dynamics within the analyzed region indicated in **a**: 3D intercellular stress field dynamics (**c**), 3D density field dynamics (**d**), 3D speed field dynamics (**e**), and 3D acceleration magnitude field dynamics (**f**). **(g**–**j)** Time series of population statistics of the intercellular stress **(g)**, cell density **(h)**, cell speed **(i)**, acceleration magnitude **(j)**. Specifically, the solid line and shaded region denote the mean and standard deviation, respectively, computed from sampled voxels within the measured 3D fields.

**Extended Data Fig. 10.**
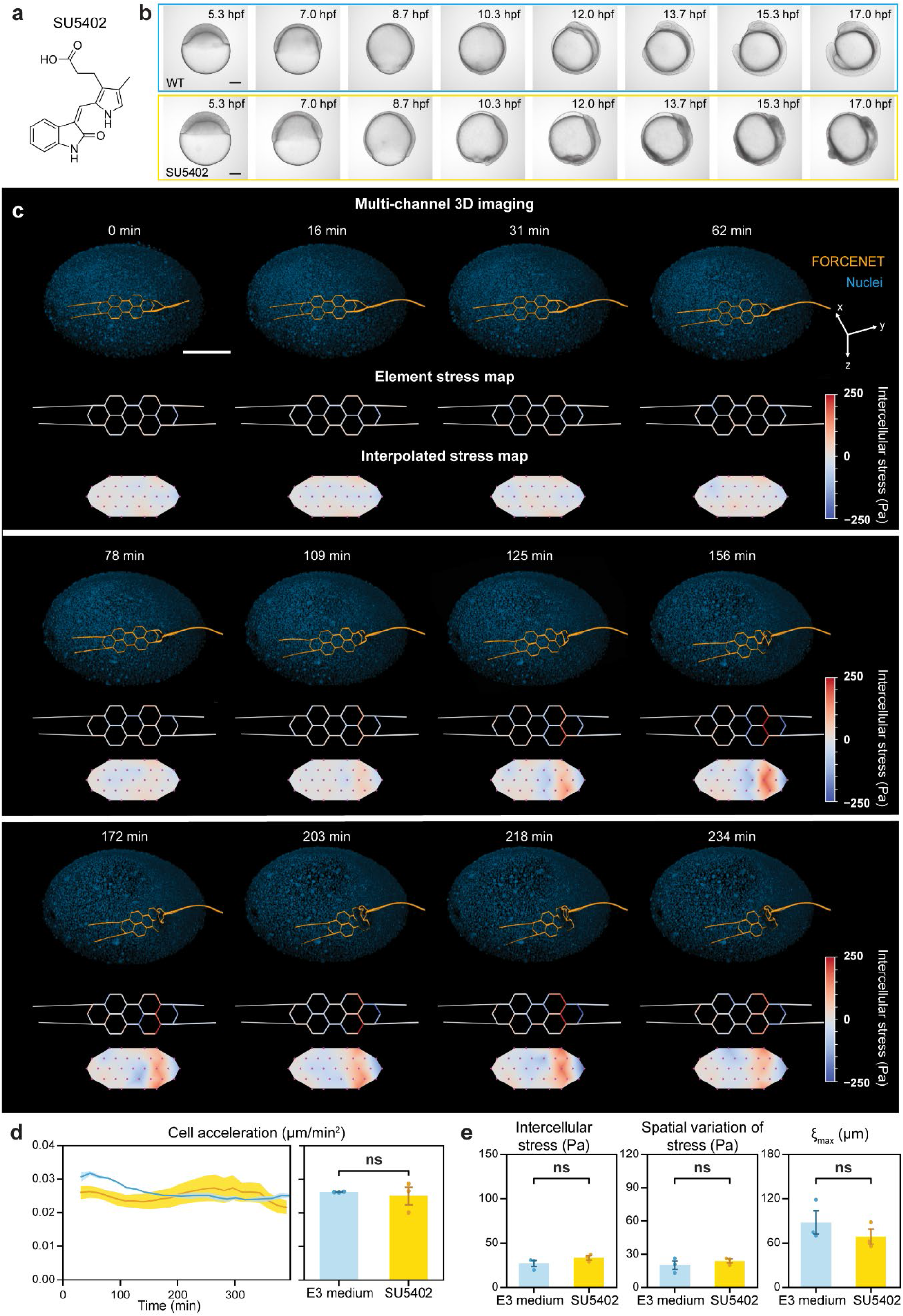
FGF signaling perturbation alters intercellular stress dynamics during zebrafish gastrulation. **a,** Chemical structure of the FGF receptor inhibitor SU5402. **b,** Time-lapse brightfield imaging of WT control (top) and SU5402-treated (bottom) zebrafish embryos. **c,** Temporal evolution of SU5402-treated embryos with implanted FORCENET, corresponding element stress maps, and interpolated intercellular stress maps. Colors in the multi-channel 3D imaging indicate nuclei (blue) and FORCENET (orange). **d,** Time series of the cell acceleration magnitude in control and SU5402-treated embryos (left) and comparison of time-averaged cell acceleration magnitude between groups (right). Lines indicate group means and shaded regions denote ± s.e.m.. Welch t-test indicated no significant difference between the two groups (p = 0.712). **e,** Time-averaged intercellular stress (left), spatial variation of stress (middle), and maximum correlation length (ξ_max_) (right) for individual embryos. (**d**–**e**) Bar values indicate the time-averaged mean, error bars indicate ± s.e.m., and individual points indicate time-averaged values per embryo. No significant differences were detected between groups (n = 3 for each group) based on a two-tailed Welch’s t-test. The p-values for intercellular stress, spatial variation of stress, and ξ_max_ were 0.218, 0.437, and 0.371, respectively.

