## Supplementary Materials for "An implantable sensor network for large-scale cellular force mapping in living embryos"

**The PDF file includes:**

Supplementary Notes 1-6

Supplementary Figure 1

Legends for Supplementary Videos 1-7

Supplementary References

**Other Supplementary Materials for this manuscript include the following:**

Supplementary Videos 1-7

### Supplementary Notes

#### Note 1: Filling ratio of different 2D lattices

The low material occupancy of FORCENET is a key design feature that minimizes tissue displacement while maintaining large-scale sensing coverage. To compare different 2D lattice geometries, we define the filling ratio, $\phi$, as the fraction of projected area occupied by polymer beams within a repeating unit cell:

$$\phi=\frac{A_{\text{beam}}}{A_{\text{total}}}$$

where $A_{\text{beam}}$ is the total projected area occupied by the beam elements within one unit cell and $A_{\text{total}}$ is the projected area of the unit cell. Throughout this analysis, each beam is approximated as a rectangle of width $w$ and length $L$, and overlap at junctions is neglected because $w \ll L$. According to crystallography, only five 2D Bravais lattice classes exist: oblique, rectangular, centered rectangular, square, and hexagonal/honeycomb (Extended Data Fig. 1b). We excluded the centered rectangular lattice from further analysis because it is not a distinct topology in terms of material occupancy. A centered rectangular lattice can be represented as an oblique lattice with an alternative choice of primitive unit cell, and therefore does not provide an independent lower bound on filling ratio beyond the oblique lattice considered below.

#### Honeycomb lattice

For a regular honeycomb lattice, the area of a unit cell is:

$$A_{\text{total}}=\frac{3\sqrt{3}}{2}L^2$$

Each of the six edges is shared by two neighboring hexagonal unit cells, giving an effective total beam length of $6L/2=3L$. Therefore,

$$A_{\text{beam}}=3wL$$

The filling ratio is

$$\phi_{\text{hex}}=\frac{2}{\sqrt{3}}\frac{w}{L}$$

For the FORCENET geometry ($w$=3 μm and $L$=30 μm)

$$\phi_{hex}=\frac{2}{\sqrt{3}}\times\frac{3}{30}\approx 11.5\%$$

#### Square lattice

For square lattice:

$$A_{\text{total}}=L^2$$

Each of the four edges is shared by two neighboring square unit cells, giving an effective total beam length of $4L/2=2L$. Therefore,

$$A_{\text{beam}}=2wL$$

The filling ratio is

$$\phi_{square}=2\frac{w}{L}$$

For the FORCENET geometry ($w$ = 3 μm and $L$ = 30 μm)

$$\phi_{\text{square}}=20\%$$

**Rectangular lattice**

For a rectangular lattice with side lengths $L_1$ and $L_2$,

$$A_{total}=L_1L_2$$

and the effective beam area per unit cell is

$$A_{\text{beam}}=w(L_1+L_2)$$

The corresponding filling ratio is

$$\phi_{\text{rect}}=\frac{w(L_1+L_2)}{L_1L_2}$$

The filling ratio is minimized when $L_1=L_2$, reducing the rectangular lattice to a square lattice. Consequently, the filling ratio of a rectangular lattice is always greater than or equal to that of a square lattice with the same characteristic length scale.

**Oblique lattice**

For an oblique lattice with side lengths $L_1$ and $L_2$ and interior angle $\theta$,

$$A_{\text{total}}=L_1L_2sin\theta$$

while

$$A_{\text{beam}}=w(L_1+L_2)$$

The filling ratio is

$$\phi_{\text{oblique}}=\frac{w(L_1+L_2)}{L_1L_2sin\theta}\geq\frac{w(L_1+L_2)}{L_1L_2}$$

with equality only when $\theta = 90^\circ$, corresponding to the rectangular lattice.

Thus, the honeycomb architecture used in FORCENET achieves approximately 42% lower material occupancy than square, rectangular, or oblique lattices of comparable dimensions. This low filling ratio enables broad spatial coverage with minimal tissue displacement, making the honeycomb geometry well suited for minimally invasive force mapping in living embryos.

**Note 2: Estimation of the bending stiffness of FORCENET force-sensing beam elements relative to zebrafish embryonic tissues**

To guide the design of FORCENET, we estimated the bending stiffness of individual force-sensing beam elements and compared it with an approximate range expected for an embryonic cell layer. The goal of this analysis was to identify a beam thickness that yields mechanical compliance on the same order of magnitude as that of surrounding tissues.

For a rectangular structure, the bending stiffness can be estimated as:

$$D = \frac{Ewh^3}{12}$$

where $E$ is Young's modulus, $w$ is the width, and $h$ is the thickness.

For FORCENET, the Young's modulus of SU-8 was measured as 4.02 GPa, the beam width is 3 μm, and the beam thickness is $h$ μm. For loading along the beam-thickness direction, the bending stiffness of the beam is therefore:

$$D_{\mathrm{device}} = \frac{(4.02\times10^{9})(3\times10^{-6})(h\times10^{-6})^{3}}{12}$$

To estimate the bending stiffness of a zebrafish embryonic cell layer, we approximated a representative cell layer as a rectangular beam with a width of 20 μm and thickness of 20 μm. Reported Young's moduli for zebrafish embryonic tissues span a broad range depending on developmental stage, tissue type, and measurement method. Previous studies have reported values of ~60 Pa for blastomere cytoplasm, ~300 Pa for the progenitor zone, and ~700 Pa for the presomitic mesoderm[1]. Other studies reported a value of ~170–200 Pa during early development[2,3]. Based on these reports, we adopted an approximate Young's modulus range of 100–700 Pa. Using this range, the estimated bending stiffness of the embryonic cell is:

$$D_{\mathrm{cell,lowest}} = \frac{(100)(20\times10^{-6})(20\times10^{-6})^{3}}{12} = 1.3\times10^{-18}\ N\cdot m^2$$

and

$$D_{\mathrm{cell,highest}} = \frac{(700)(20\times10^{-6})(20\times10^{-6})^{3}}{12} = 9.3\times10^{-18}\ N\cdot m^2$$

Comparing these estimates with the calculated bending stiffness of FORCENET beams indicates that beam thicknesses near 200 nm produce bending stiffness within the same order of magnitude as that estimated for a single-cell-thick embryonic tissue layer (Fig. 1f). Although this simplified analysis does not capture the complex geometry and viscoelastic behavior of living cells, it provides a useful design criterion for minimizing the mechanical mismatch between FORCENET and the surrounding tissues.

**Note 3: Comparison of bending stiffness between the insertion needle and force-sensing beam**

Because both the insertion needle and the individual force-sensing beam of FORCENET are made from the same material, the ratio of bending stiffness is determined solely by their geometries. For a rectangular beam, the second moment of area for bending about the width axis is:

$$I = \frac{wh^3}{12}$$

where $w$ is the beam width and $h$ is the beam thickness in the bending direction.

For the insertion needle:

$$w_{\text{needle}} = 8\ \mu m$$

$$h_{\text{needle}} = 10\ \mu m$$

For an individual force-sensing beam:

$$w_{\text{beam}} = 3\ \mu m$$

$$h_{\text{beam}} = 0.18\ \mu m$$

Thus, the ratio of bending stiffness between the insertion needle and the force-sensing beam is:

$$\frac{8 \times 10^3}{3 \times 0.18^3} \approx 4.6 \times 10^5$$

Thus, the insertion needle is approximately $4.6\times10^5$ times stiffer than an individual force-sensing beam. Consequently, the needle provides sufficient rigidity for manipulation and implantation, whereas the ultrathin FORCENET beams remain highly compliant and mechanically sensitive to forces generated by surrounding cells.

**Note 4: Estimation of stress-sensing uncertainty**

We estimated the stress-sensing uncertainty of FORCENET using a cantilever model of a single force-sensing beam element, with one end fixed and the beam subjected to a uniform load along the beam thickness direction (Extended Data Fig. 1d). We focused on two dominant experimental sources of uncertainty: variation in beam thickness and uncertainty in node detection.

For a cantilever beam of length $L$, width $w$, thickness $h$, and Young' modulus $E$, the relationship between the applied uniform stress $\sigma$ and the end-node displacement $\delta$ (Extended Data Fig. 1d) is:

$$\sigma = \frac{2Eh^3}{3L^4}\delta$$

Because the uncertainties in beam thickness and nodal localization are independent, the propagated uncertainty can be approximated as:

$$\Delta\sigma = \sqrt{\left(\frac{\partial\sigma}{\partial h}\Delta h\right)^2 + \left(\frac{\partial\sigma}{\partial\delta}\Delta\delta\right)^2} = \sqrt{\left(3\sigma\frac{\Delta h}{h}\right)^2 + \left(\sigma\frac{\Delta\delta}{\delta}\right)^2}$$

Therefore, the relative stress uncertainty is:

$$\frac{\Delta\sigma}{\sigma} = \sqrt{\left(3\frac{\Delta h}{h}\right)^2 + \left(\frac{\Delta\delta}{\delta}\right)^2}$$

Experimentally, the beam thickness measure across a whole wafer was 182 ± 2 nm (mean ± s.d.; Extended Data Fig. 2d), giving a thickness-contributed relative uncertainty of

$$3\frac{\Delta h}{h} \sim 3.3\%$$

To estimate the contribution from node detection uncertainty, we considered a representative stress of 50 Pa, which is towards the lower end of the measured intercellular stress range, corresponding to higher end of the relative uncertainty. For this stress level,

$$\delta = \frac{3L^4}{2Eh^3}\sigma = 2.6\ \mu m$$

The nodal translation $\delta$ was obtained from node localization in the reconstructed FORCENET geometry (Extended Data Fig. 6b). The localization precision can be estimated using the Cramér–Rao bound commonly used in single-molecule localization microscopy[4] as

$$\Delta\delta \sim \frac{\text{imaging resolution}}{\sqrt{N_{\text{photon}}}}$$

where $N_{\text{photon}}$ is the number of detected photons within a diffraction-limited spot. Under our imaging condition, we typically detected approximately 100-1,000 photons. For two-photon excitation at 920 nm with a 0.95 NA objective, the lateral resolution is approximately 300-400 nm and the axial resolution is approximately 1.2-1.8 µm. Using the most conservative imaging resolution estimate of 1.8 μm and $N_{\text{photon}}$=100, we obtained $\Delta\delta$~180 nm. Thus the node-detection-contributed relative uncertainty is

$$\frac{\Delta\delta}{\delta} \sim 6.9\%$$

Combining the thickness and localization contributions gives an estimated relative stress uncertainty of

$$\frac{\Delta\sigma}{\sigma} \sim \sqrt{(3.3\%)^2 + (6.9\%)^2} = 7.6\%$$

**Note 5: Characterization of FORCENET stress reconstruction**

FORCENET reconstructs an averaged intercellular stress along the thickness directions of individual force-sensing beam elements. This finite spatial averaging defines a practical spatial resolution for the stress reconstruction. To characterize this resolution, we performed the reconstruction pipeline under applied stress gratings with controlled spatial length scales and orientations (Extended Data Fig. 7):

$$\sigma_X(x, y) = \sigma_A\, cos[\frac{\pi(x - x_c)}{L_{\text{peak-valley}}}]$$

and

$$\sigma_Y(x, y) = \sigma_A\, cos[\frac{\pi(y - y_c)}{L_{\text{peak-valley}}}]$$

where $\sigma_A$ is the stress amplitude, $L_{\text{peak-valley}}$ is the physical distance between a local stress maximum and the adjacent minimum, and $(x_c, y_c)$ is the geometric center of FORCENET. The phase of each stress grating was chosen such that a stress maximum was centered on the sensing region. This centering minimized loading-position differences across gratings and allowed the effect of spatial length scale to be evaluated more directly. In this analysis, $\sigma_A$ was held fixed at 60 Pa, and $L_{\text{peak-valley}}$ was varied from 20 to 160 μm, spanning length scales from below the beam-element length to a substantial fraction of the full FORCENET sensing region.

For the forward simulation, each stress grating was applied to the FORCENET finite element model as a spatially varying distributed load along the beam thickness direction. The applied stress field was sampled along the centerline of each force-sensing beam element and converted into piecewise-linear distributed loads. The ends of the interconnecting ribbons outside the force-sensing region were fixed in all translational and rotational degrees of freedom, approximating the experimental boundary condition. Forward finite element analysis was then used to calculate the resulting 3D deformation of FORCENET.

For reconstruction, the applied stress grating was not provided to the reconstruction pipeline. Instead, the 3D coordinates of the deformed FORCENET nodes and internodes were extracted from the forward simulation and used as the only input to the reconstruction. The reconstruction pipeline calculated nodal forces from the simulated deformation and converted these nodal forces into stresses on individual force-sensing beam elements using the same procedure applied to experimental data.

For comparison, the ground-truth element stress map was computed by averaging the applied stress gratings over each beam element. The ground truth and reconstructed element stress maps were then interpolated using the barycentric interpolation procedure used for experimental stress maps (Methods).

**Note 6: Spatial autocorrelation and cross-correlation analysis**

For each time point $t$, spatial autocorrelation and cross-correlation analyses were performed on 3D grids $\boldsymbol{x} = (x, y, z) \in \mathbb{Z}^3$. Correlations were computed after subtracting the mean value of each field within its valid masked region.

For the intercellular stress autocorrelation, the valid mask corresponded to the interpolated stress field defined by the FORCENET (Fig. 3h). For the cross-correlation analysis, the valid mask corresponded to the device segmentation dilated by approximately 20–30 μm from the device centerline, as illustrated by the dashed box in Fig. 4a.

The raw cross-correlation between two fields at a spatial displacement $\boldsymbol{\Delta}$ was defined as

$$C_{AB,t}^{\mathrm{raw}}(\boldsymbol{\Delta}) = \sum_{x} A'_t(\boldsymbol{x})\ B'_t(\boldsymbol{x} + \boldsymbol{\Delta})$$

where $A'_t$ and $B'_t$ denote the masked, mean-centered fields at time $t$.

Because finite image volumes contain progressively fewer overlapping voxels at larger spatial displacements, the raw cross-correlation was corrected by the overlap of the corresponding binary validity masks,

$$N_{AB,t}(\boldsymbol{\Delta}) = \sum_{x} M_{A,t}(\boldsymbol{x})\ M_{B,t}(\boldsymbol{x} + \boldsymbol{\Delta})$$

where $M_{A,t}$ and $M_{B,t}$ are binary masks indicating valid voxels. The overlap-corrected correlation was then computed as

$$\widetilde{C_{AB,t}}(\Delta) = \frac{C_{AB,t}^{\text{raw}}(\mathbf{\Delta})}{N_{AB,t}(\mathbf{\Delta}) + \varepsilon_t}$$

where $\varepsilon_t$ was set to 5% of the maximum value of $N_{AB,t}$ to ensure the numerical stability at displacements with very small overlapping voxels.

To facilitate comparison between different variable pairs, the overlap-corrected correlation was further normalized by the standard deviations of the two fields,

$$\Gamma_{AB,t}(\mathbf{\Delta}) = \frac{\widetilde{C_{AB,t}}(\mathbf{\Delta})}{\sigma_{A,t}\sigma_{B,t}}$$

where $\sigma_{A,t}$ and $\sigma_{B,t}$ denote the standard deviations (s.d.) of the masked fields. This normalization yields a dimensionless correlation metric that is independent of the absolute variance of the underlying measurements.

To obtain the radially averaged cross-correlation functions, the 3D normalized correlation function was averaged over all displacement vectors with magnitude $|\mathbf{\Delta}|$ falling within a radial bin centered at distance $r$.

$$g_{AB,t}(r) =< \Gamma_{AB,t}(\mathbf{\Delta}) >_{|\Delta|\approx \mathrm{r}}$$

Spatial autocorrelation was computed as the special case $A = B$.

**Supplementary Figures**

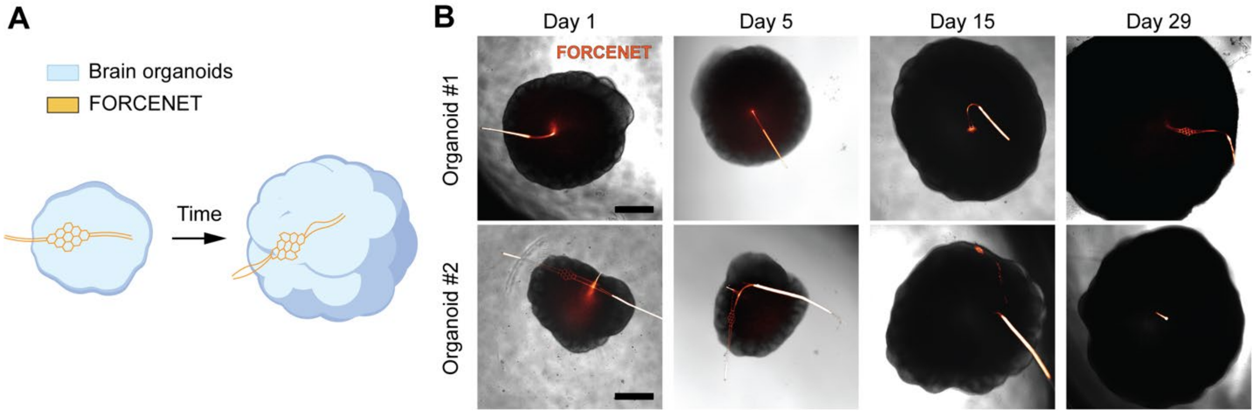


**Supplementary Fig. 1 | FORCENET integrates with developing human cortical organoids. a,** Schematic of a FORCENET integrated within a developing human cortical organoid. **b,** Representative correlative brightfield-fluorescence images of two FORCENET (orange)-integrated human cortical organoids cultured for 29 days post FORCENET integration. FORCENET was integrated at Day 30 post organoids formation. Scale bars, 500 μm.

### Legends for Supplementary Videos

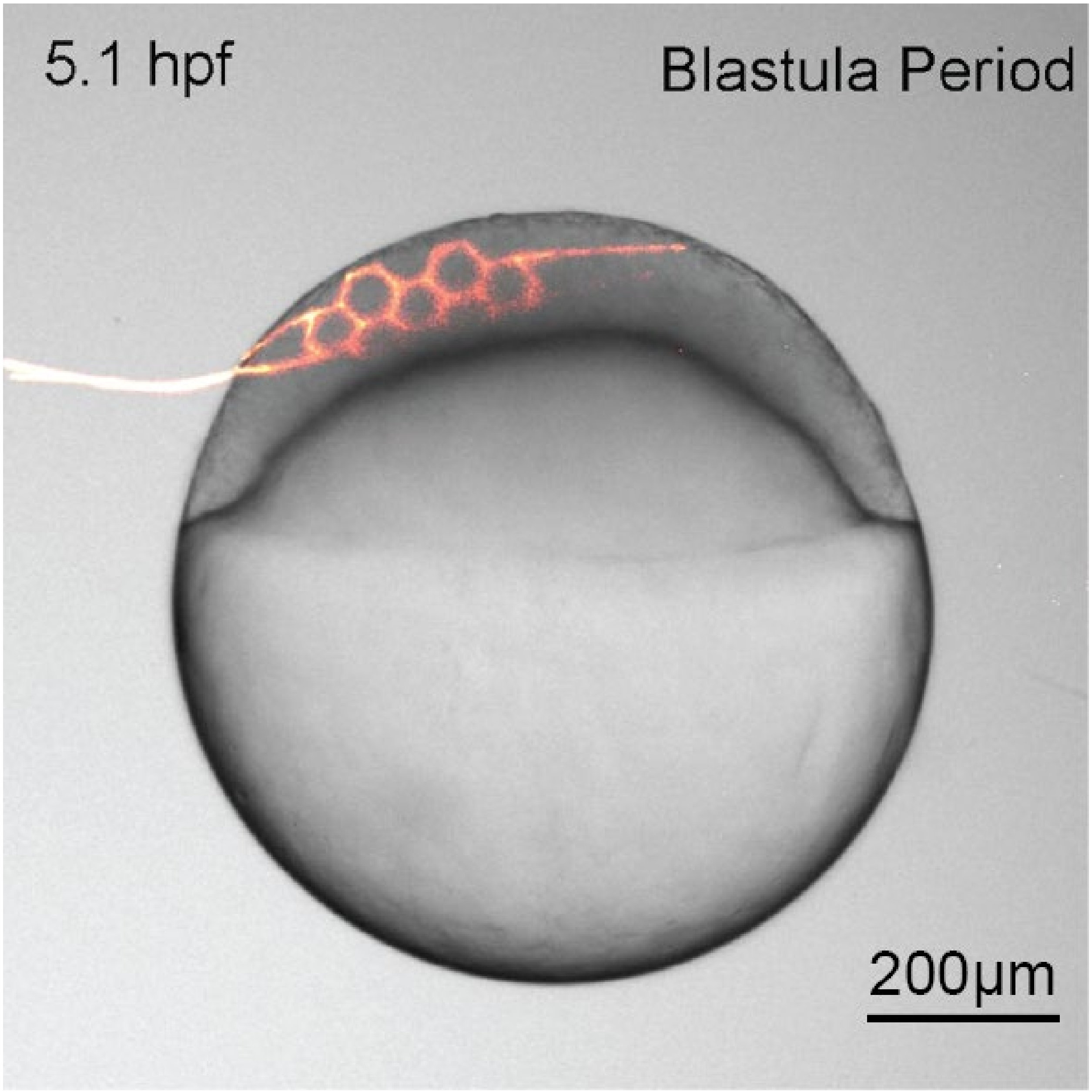


**Supplementary Video 1.** Correlative brightfield-fluorescence time-lapse imaging of zebrafish development following FORCENET (orange) implantation. Scale bar, 200 µm.

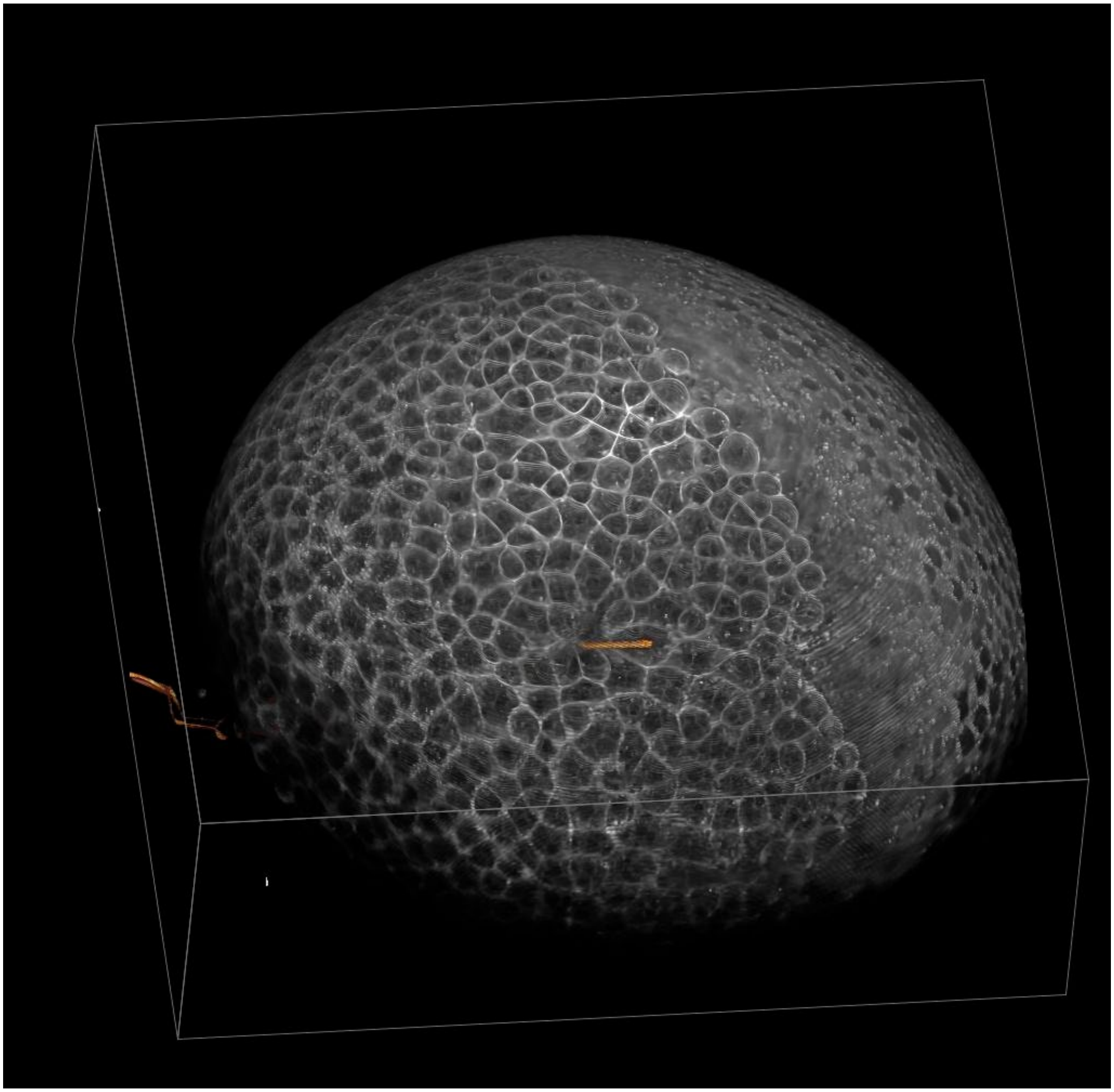

**Supplementary Video 2.** A fly-through of two-photon z-stacks showing the interface between FORCENET (orange) and cell membranes (white). Bounding box dimension: 201 × 766 × 766 $\mu m^3$ (z × y × x).

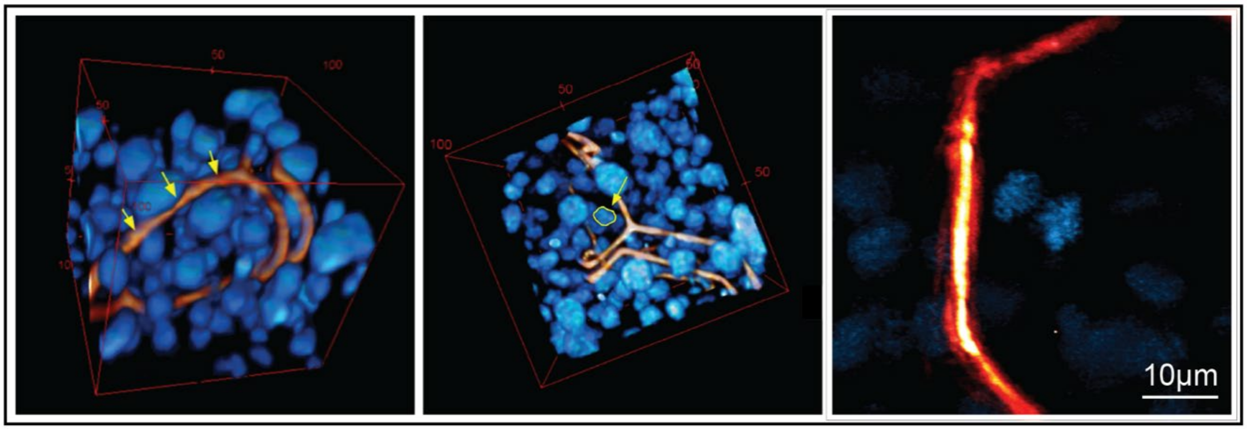


**Supplementary Video 3.** Video of the FORCENET–tissue interface. FORCENET is shown in orange, and nuclei are shown in blue. The movie shows three examples: (1) cellular movements deform FORCENET beams, (2) cells pass through FORCENET, and (3) cell mitosis occurs near FORCENET. The first two segments were captured using two-photon microscopy, while the final segment is a maximum-intensity projection from lattice light-sheet microscopy. For the three parts of the movie, bounding box dimensions: 84 × 84 × 82 μm³ for (1); 92 × 97 × 90 μm³ for (2); and scale bar: 30 μm for (3).

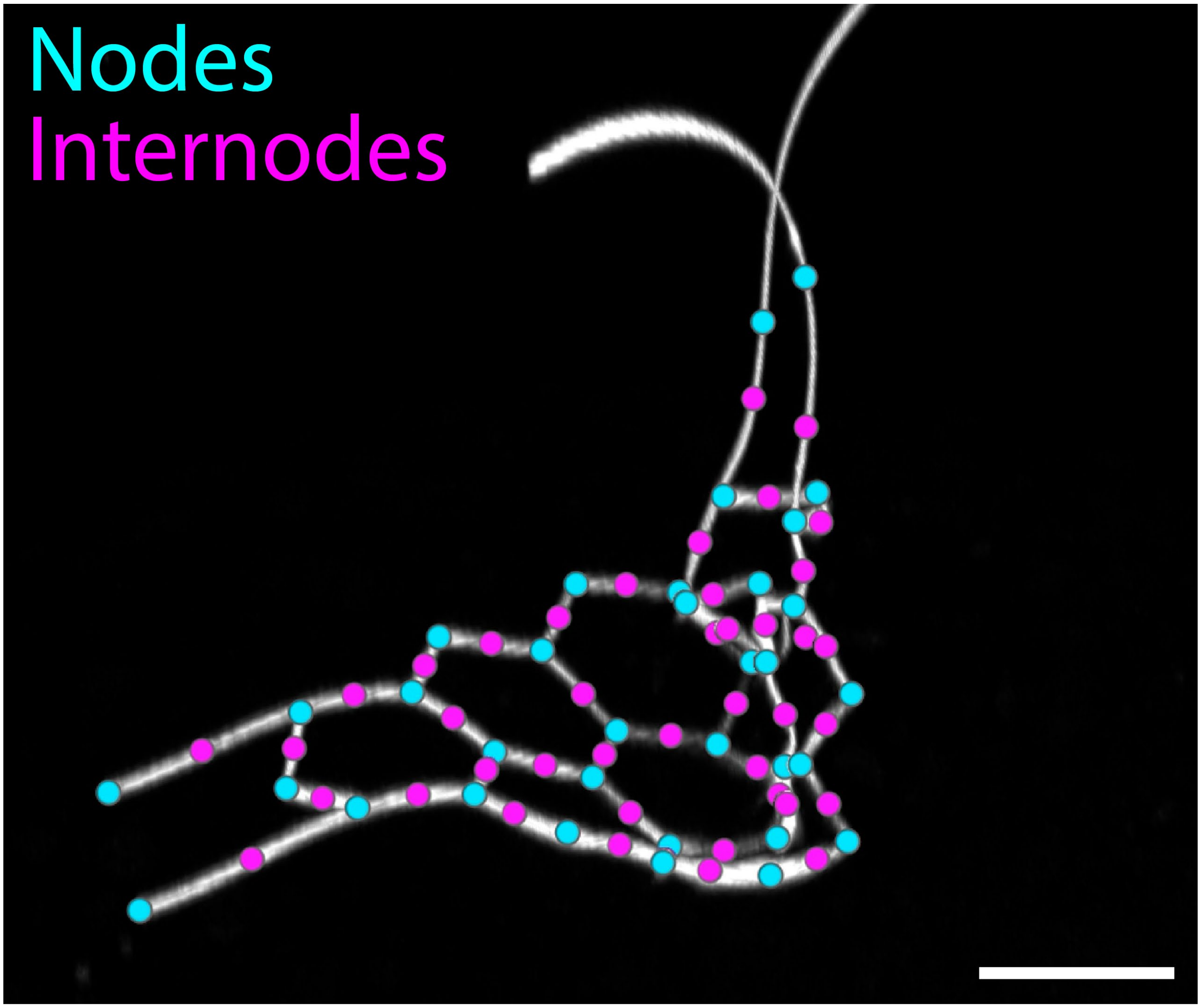


**Supplementary Video 4.** Overview of the computational analysis pipeline showing how raw two-photon images of FORCENET (white) are processed to extract nodal and internodal positions, from which device deformation and stresses are determined. Lastly, the result of the computational pipeline is demonstrated wherein the multi-channel imaging, device segmentation, and the resulting element stress map and interpolated stress map are demonstrated. All scale bars: 100 µm.

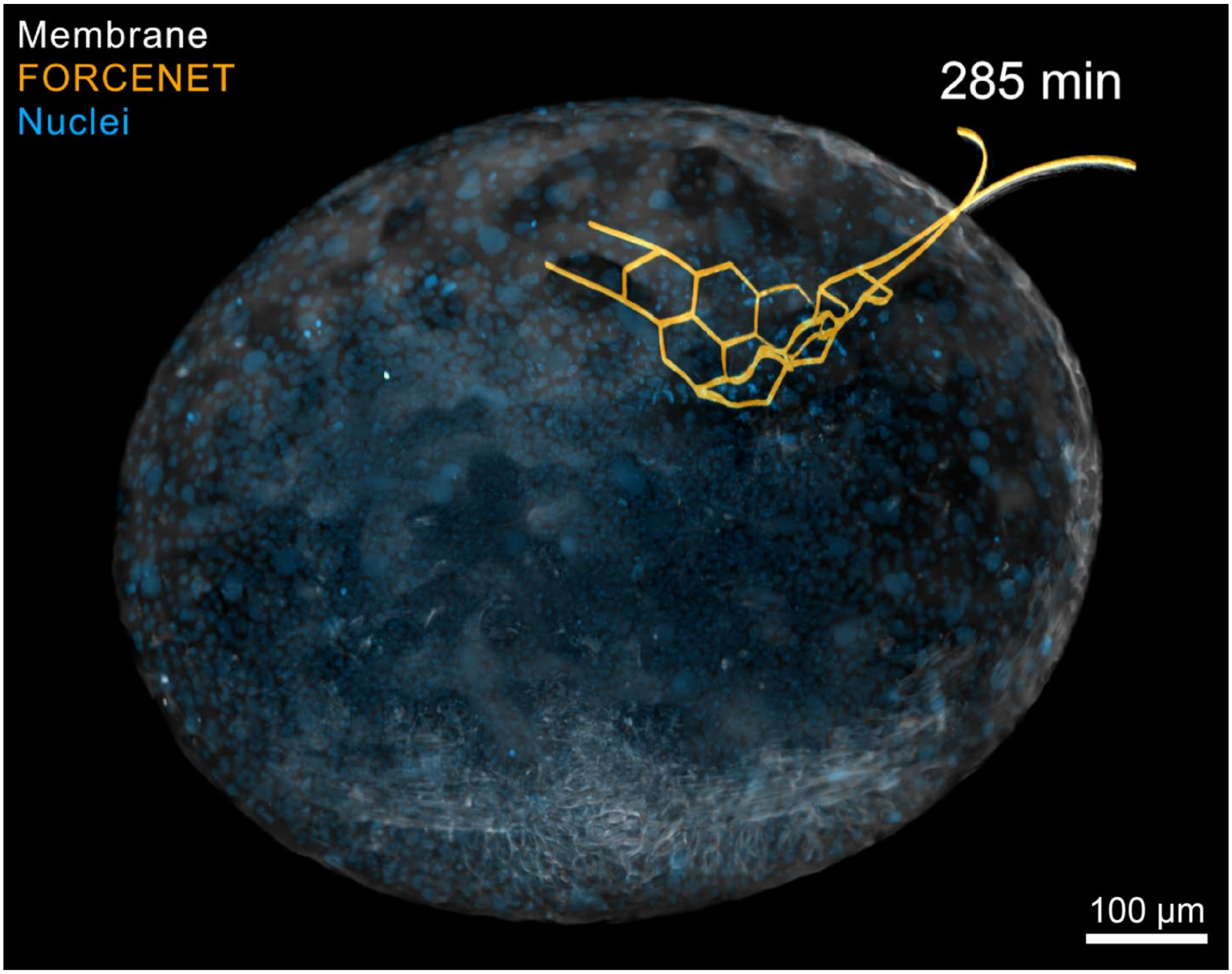


**Supplementary Video 5.** FORCENET maps intercellular stress across zebrafish gastrulation. Images were acquired using two-photon microscopy at 15.8 min intervals. Scale bar, 100 µm. The movie cycles through a fly-through of the imaging z-stacks showing the FORCENET–embryo interface; visualization of the raw multichannel data, including nuclei (blue), FORCENET (orange), and membranes (white); and a multi-panel time-series video of the device deformation, element stress maps, and interpolated stress maps. The final segment shows the propagation of stress across FORCENET. In the multi-panel video, all scale bars: 50 µm.

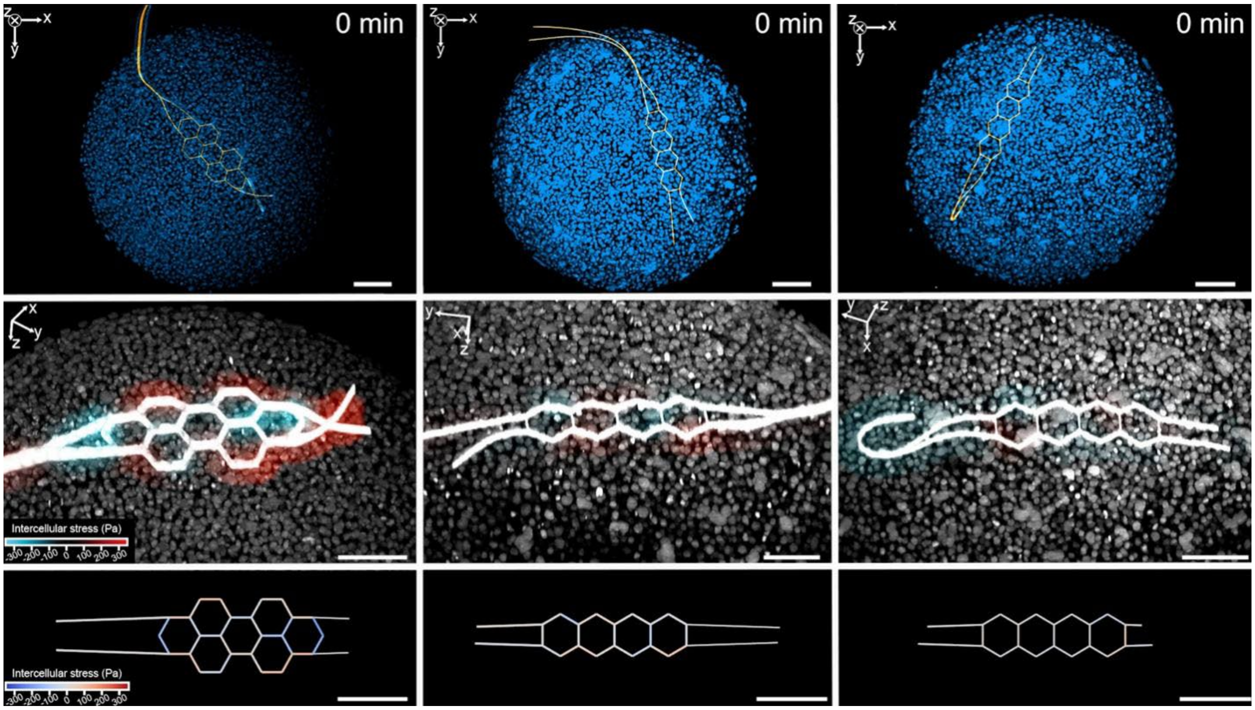


**Supplementary Video 6.** FORCENET measures intercellular stress across multiple embryos. Embryo #1 (left) was imaged for 25 time points at 15.8 min intervals, whereas Embryos #2 (middle) and #3 (right) were imaged for 35 time points at 15.6 min intervals. The top row shows the raw multichannel data, with nuclei in blue, and FORCENET in orange. The middle row shows the FORCENET and nuclei channels, both in white, overlaid with the interpolated 3D stress field. The bottom row shows the element stress maps. In the middle and bottom rows, colormaps denote signed intercellular stress values. All scale bars: 100 μm.

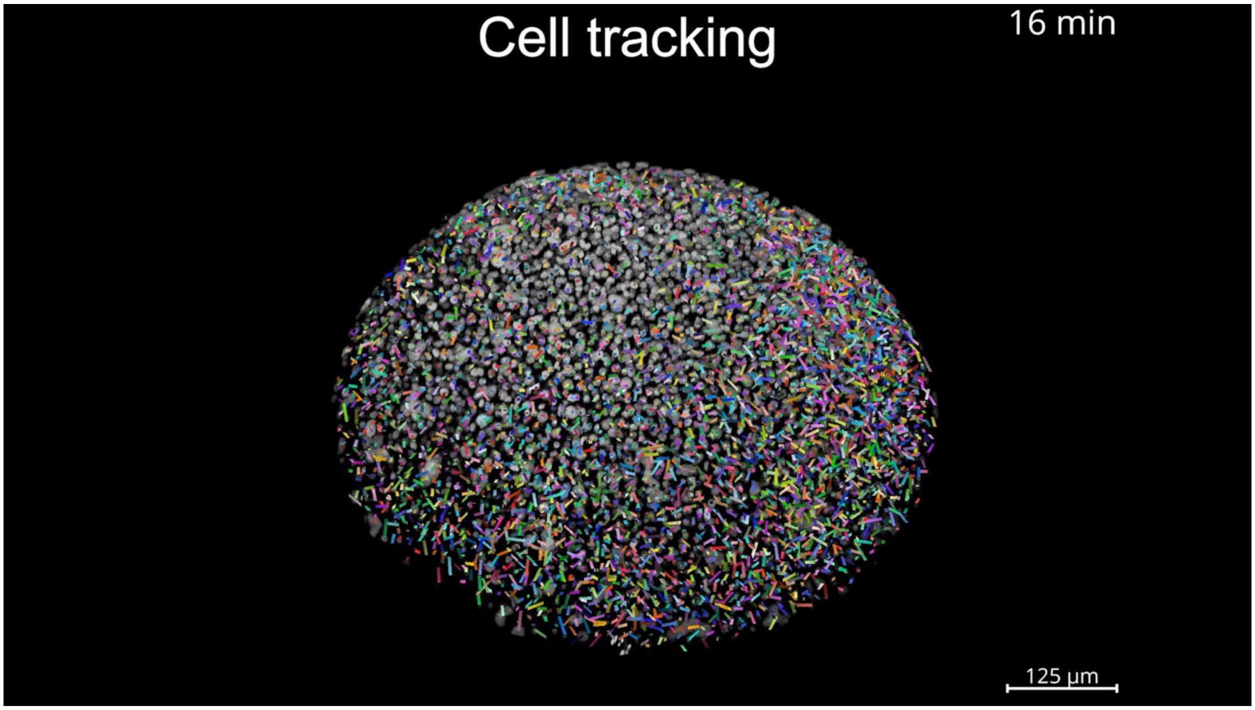


**Supplementary Video 7.** 3D cell segmentation and tracking using the nuclear channel (white) from the 3D multi-channel imaging. The movie then displays the temporal evolution of the intercellular stress field alongside the cell density, speed, and acceleration magnitude fields. Bounding box: $75 \times 427 \times 190$ $\mu m^3$ ($z \times y \times x$).
